# Sexual display behaviour follows consistent sex-specific reaction norms across latitude in response to operational sex-ratio

**DOI:** 10.1101/2024.05.08.592656

**Authors:** Ivain Martinossi-Allibert, Sebastian Wacker, Claudia Aparicio Estalella, Charlotta Kvarnemo, Trond Amundsen

## Abstract

Predicting the strength and direction of sexual selection is a challenging task for evolutionary theory, as the effects of ecological factors, social environment, and behavioural plasticity, all need to be taken into account. The Operational Sex Ratio (OSR) is a key variable, which has been shown to (i) affect the strength and direction of mating competition, as a social environment cue, and (ii) be affected itself by ecological conditions through sex-specific environmental effects. Yet, gaining a global view of (i) and (ii) in wild populations represents an arduous but necessary step to further our understanding of sexual selection dynamics in the wild. Here, we address this challenge by using reaction norms. We conducted an extensive field study on the two-spotted goby *Pomatoschistus flavescens*, monitoring six populations along a latitudinal gradient during an entire breeding season. Doing so, we compared across populations the temporal trajectories in social environment and sexual displays, which is unprecedented. We develop a reaction norm framework based on OSR theory to analyse the data. We show that what appears to be tremendous variation in sexual displays across populations and sampling times, follows consistent rules: sexual display behaviour follows behavioural reaction norms in response to the social environment that are consistent across populations, but social environment fluctuations are specific to each population. Recording behaviour not only over time, but also along a latitudinal gradient where ecological conditions change and in turn affect OSR, was necessary to gain insight into the relationship between social environment and sexual displays, which in turn contributes to sexual selection dynamics.

## Introduction

From an individual’s perspective, the decision to engage in mating is probably the most important turning point of a life history trajectory. Accordingly, reproductive strategies are finely tuned to external cues for resources, predation, as well as potential mates and competitors (*i*.*e*., the social environment). Population-level processes of sexual selection emerge from these individual decisions, and thus, understanding the patterns in strength and direction of sexual selection that shape life and its diversity requires an understanding of what drives individual reproductive strategies in a population context.

To that end, we distinguish two types of drivers likely to shape individual reproductive strategies and reproductive behaviours. First, predictable environmental cycles, such as those driven by seasonality, are central in determining reproductive strategies (Conover 1992). For example, in many species individuals strive to time the birth of their young with a predictable resource peak (Martin 1987, Rutberg 1987). Classically, environmental cycles are expected to shape consistent reproductive strategies, even though these are currently being disrupted by rapid and pervasive environmental change (*e*.*g*., Asch et al. 2019, Visser and Gienapp 2019). Second, information on conspecifics, sometimes also referred to as “social environment”, shapes reproductive strategies and associated behaviours; conspecifics are both potential mates and competitors, and their abundance in the mating pool affects the optimal strategy of a given individual (e.g. Formica and Tuttle 2009, Webster et al.2010). Adapting a strategy to the social environment can lead to complex and unstable frequency dependent dynamics that require individuals to respond to rapidly changing conditions, or in other words to be behaviourally plastic (Dugatkin and Reeve 1998, Piersma and Drent 2003, Dubois et al. 2010). To evaluate how sexual selection operates in an ecological context, it is thus important to understand the impact of fluctuations in the social environment, as well as link them to their ecological drivers.

The Operational Sex Ratio (OSR) framework is useful in this context. It proposes that the relative abundances of males and females in the mating pool (the OSR) predicts the direction and intensity of mating competition, and thus the frequency of associated behaviours (Emlen and Oring 1977, Kvarnemo and Ahnesjö 1996, 2002). A straightforward link between OSR and competition relies on the assumption that it is adaptive to compete whenever competitors are abundant. However, that may not be the case, for example when alternative reproductive tactics are available (Jeffery et al. 2018, Heimerl et al. 2022). Nevertheless, assuming that competition is the best strategy, the OSR framework predicts that individuals of the most abundant sex in the mating pool should compete and court more intensely, reflecting stronger sexual selection on that sex.

The OSR is likely to vary across species, across populations, and even within populations over time (Ahnesjö et al. 2008; Miller & Svensson 2014). These variations over different scales occur because females and males have different optimal reproductive strategies implying for example different growth rates, survival rates, timing of sexual maturity, and potential reproductive rates (PRR, Clutton-Brock and Parker 1992), but also different sensitivities to environmental fluctuations. When the sexes respond differently to environmental cues, it may also lead to mismatches in the timing of reproduction and fluctuations in mating pool composition (e.g. Forsgren et al. 2004, Weladji et al. 2017).

If the OSR is consistently biased over evolutionary time scales, adaptive evolution of behaviours is expected, such as courtship of the rarer sex evolving in the more abundant one (Kokko et al. 2012). For example, in the pipefish *Syngnathus typhle*, potential reproductive rate is consistently limited by the brood pouch of the male, a parental care feature that has evolved in pipefish and seahorses, and leads to frequent female courtship and female-female competition (Berglund 1994). In many other species, it is the fecundity of the female or female-biased parental care that constrains reproductive output and leads to male-male competition (Kvarnemo & Ahnesjö 1996, Janicke et al. 2016). The role of OSR in driving mating competition on such a scale has gathered support from meta-analyses (Weir et al. 2011, Janicke and Morrow 2018), and from experimental evolution (Booksmythe et al. 2014), although the meta-analyses suggest a relatively weak explanatory power. This weak power is not surprising considering that many other factors are likely to affect the strength of sexual selection (Klug et al. 2010), for example the existence of alternative reproductive tactics (Jeffery et al. 2018) or the mode of parental care, which both can be ecologically and phylogenetically constrained.

In most populations it is also expected that OSR fluctuates on a short time scale, due to its dependence on ecological factors with sex-specific effects (Emlen and Oring 1977, Ahensjö et al. 2008, Kvarnemo & Simmons 2013). For example, extrinsic mortality (Forsgren et al. 2004), availability of resources required for mating (Garcìa-Berro et al. 2019), extreme events such as heat waves (Garcia-Roa et al. 2020) or availability of food (Gwynne and Simmons 1990) can affect the sexes differently, leading to OSR fluctuations.

In the case of short-term fluctuations, OSR theory assumes that individuals plastically adjust their behaviour to the perceived social environment. Empirical work generally supports plastic response of sexual display to OSR (reviewed in de Jong et al. 2012, see also: Chuard et al. 2016, Weladji et al. 2017, Villarreal et al. 2018, Munõz-Arroyo et al. 2019, Driscoll et al. 2022, Chuard et al. 2022). However, as pointed out first by de Jong et al. (2012), these studies lack a consistent and explicit framework, which makes it difficult to draw comparisons and generalise. For example, although most observations are made in controlled environments, there is no consistency in methods to control for encounter frequencies between individuals, and the expected relationship between the behaviour performed and the social environment is not always explicit.

Here, we propose a simple framework to test predictions of OSR theory in wild populations. This framework deals with the complexity of distinguishing Adult Sex Ratio (ASR) and OSR (Carmona-Isunza et al. 2017), which is crucial in the field as all adults present in a population may not be qualified to mate at a given time. The framework allows an evaluation of propensities to perform focal behaviours estimated from field observations, by correcting for encounter rates of the different classes of individuals. We describe the relationship between behaviour propensity and social environment as a reaction norm, which is a familiar concept to most in the fields of evolutionary ecology (Dingemanse et al. 2010).

We use this framework to analyse data from a large-scale field study of sexual diplay in the two-spotted goby *Pomatoschistus flavescens*, surveying six wild populations spread across a latitude gradient of two thousands kilometers along the Norwegian coastline, during an entire breeding season (spring and summer 2022). Sexual displays consist of courtship initiation and of same-sex agonistic interactions performed by both sexes. This study is unprecedented in allowing us to (i) compare the simultaneous temporal trajectories of sexual display behaviours in multiple populations across an environmental gradient, and (ii) assess how social environment drives sexual display behaviours in wild populations on such a spatial and temporal scale. We discuss how ecological factors affect the social environment, with cascading effects on sexual display behaviour.

## Materials and Methods

### 1. Study species and reproductive behaviour

The two-spotted goby Pomatoschistus flavescens is a small semi-pelagic fish widely distributed in the Northeast Atlantic. It is usually considered to have an annual life cycle, and in Norway the breeding period spans from March to August, concluding with the mortality of the spawners (Skolbekken et al. 2001). It usually breeds within the littoral zone, typically in shallow waters of up to 5 meters deep and in kelp habitats where males can hold a nest. As a consequence, temperature fluctuations within this habitat can vary significantly from year to year throughout the mating season and even from day to day due to weather conditions. Both sexes have distinct sexually selected ornaments and may perform courtship displays and same-sex agonistic behaviors, making this species particularly suited to the study of rapid variation in sexual display behaviour (Amundsen 2018). Male ornamentation consists of large and pigmented dorsal and anal fins (banded red/purple and dark, respectively), which can be sometimes accompanied by a general reddish pigmentation of the body and in particular of the head (Wacker et al. 2013). Males perform parental care usually for the clutches of several females, until hatching of the eggs, and easily breed in artificial nests placed in the wild (Mobley et al. 2009, Amundsen 2018). Small males occasionally engage in alternative reproductive tactics by sneaking into a nest during fertilisation (Utne-Palm et al. 2015). Male courtship starts with a *fin display*, which consists in extending the dorsal and anal fins. This display can also be directed at males in antagonistic interactions, and in this case can be followed by a *chase*, which consists of brief pursuits between males with rapid approach movements, to our knowledge never leading to actual biting.

Female ornamentation during the breeding season is characterised by a belly full of bright orange eggs (Amundsen and Forsgren 2001), complemented by carotenoid-based orange pigmentation of the skin (Svensson et al. 2006). This ornament is thought to advertise receptivity, and perhaps to also be a signal of fecundity (Svensson et al. 2006). Females can further highlight their orange belly and eggs by turning the rest of their body almost transparent (Sköld et al. 2008). We refer to this colour change as *glowing* behaviour. In addition, females perform an arching of their body that highlights the roundness of their egg-filled belly that is often directed towards males during courtship, but also towards females in antagonistic interactions (Amundsen and Forsgren 2001). We refer to this behaviour as the *hook* behaviour, because the body posture resembles a hook in shape.

In male-female courtship, if a female responds positively to a fin display (by either approaching the male or performing a hook or a glow), the courtship may continue by the male performing a characteristic undulating lead swim towards his nest.

### 2. Field data collection

#### Study populations

The six study sites extend from the western coast of Sweden in the South (58.2º N) to northern Norway (68.6º N, Figure 1). In Supplementary File F1, we present detailed maps showing the location of each population, each sub-location and the position of the observation transects. The populations were chosen to cover the latitude range with regularly spaced sampling sites, while favouring sites where the species had been observed in abundance in the past (personal observations) to ensure that suitable habitat and sufficient population density could be found.

**Figure 1.**
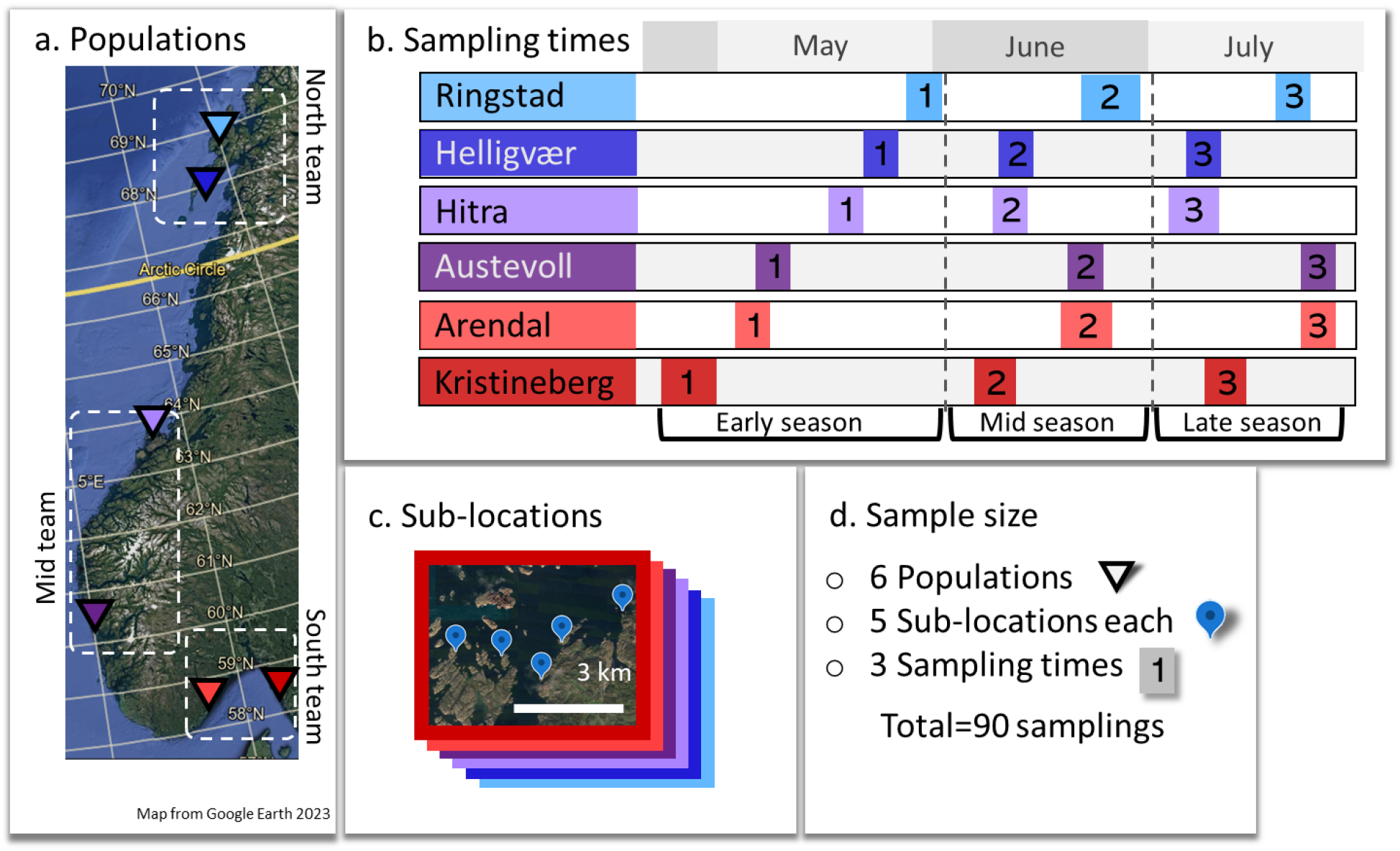
Sampling scheme for behaviour and population census of the two-spotted goby (Pomatoschistus flavescens). The locations of the 6 populations are marked on the map(a), together with the time of sampling in early, mid- and late season for each (b). For each population, 5 replicate sublocations (c) were sampled at each time period, for a total of 30 sublocations and 90 samplings (d). Three field teams covered two of the 6 populations each. Detailed locations and sampling date of each sampling site and transect are given in supplementary figure S1, and supplementary file F1.

#### Sampling design

The sampling design is summarised in Figure 1. For each of the 6 study populations, 5 sub-locations were chosen, representing replicated measurement of the populations (see Supplementary File F1 for exact locations). In each of the 30 sub-locations, we recorded behaviour and population censuses three times over the course of the breeding season (from April to July 2022), representing early, mid- and late-season observations. The sampling started later in northern locations, because the breeding season was expected to be delayed due slower water temperature rise and delayed primary productivity. The sampling was executed by three different field teams, each in charge of 2 of the 6 populations (Figure 1.a). The three team leaders trained together prior to the start of the field season, in order to standardise methods and ensure a minimal bias due to differences in measurement or observation methods. Before undertaking transect surveys in the water, the exact starting and ending points of the transects had been determined to ensure they would consistently occur in the same space, thereby facilitating the collection of consistent information regarding changes throughout the season. The three team leaders recorded all the behaviour observations and population censuses throughout the study, and were supported by other team members for less observer-bias prone tasks.

#### Behavioural transects

Recording of behaviour was done through underwater visual censuses done by one snorkelling observer swimming along the shore, using a standardised form (supplementary S2). Transects were established in suitable breeding habitat (abundant kelp habitat close to the shore) and the same transects were sampled repeatedly throughout the breeding season. Transects were approximately 1m wide and length varied depending on the disposition of the suitable breeding habitat patch (average length 120m). In short, the behaviour form recorded numbers of behaviour observed, for both courtship initiation and antagonistic behaviours. For courtship, if the observer witnessed the beginning of a courtship interaction, the observer noted which sex initiated the behaviour (female initiated, male initiated) and how the other individual responded. If the observer witnessed ongoing courtship, it was recorded as “mutual courtship” (initiator unknown). Each transect was swum twice consecutively (one time in each direction). When analysing the data, we distinguished two main types of behaviour: courtship initiation and same-sex agonistic behaviours. Courtship initiation from male towards female includes *fin display*, and *fin display* followed by *lead swim*. Courtship initiation from female towards male includes the *hook* and *glow* behaviours. Same-sex agonistic behaviours consist of *fin display* from male to male, combined with the eventual follow up of a *chase*, and *hook* and *glow* from female to female, grouped together. To ensure that there was minimal alteration of individual behaviour due to human presence, data collection of behaviour was the first task conducted during a field day (prior to captures for phenotype measurements).

#### Population census and social environment

The same transect as for the behavioural recording was swum by a snorkelling observer twice (one time in each direction) to count the number of individuals present. This was done in alternance to the behavioural observations (also one time in each direction). Individuals were sexed, meaning that they were attributed to one of three categories: female, male or non-sexually mature adult. In addition, females were attributed to one of three roundness categories representing the stage of egg maturation:

1. females showing no roundness or very little, deemed not ready to spawn
2. females showing substantial roundness, could be close to spawning
3. females showing maximum roundness, ready to spawn

Supplementary Figure S3 shows example pictures of females attributed to different roundness classes. We assume that all females of class 2 and 3 were in the mating pool, and that all males observed in the breeding habitat were in the mating pool. This could be an overestimation of male presence in the mating pool, however it seems reasonable given the biology of the species: most individuals have only 1 season to reproduce, so they are unlikely to postpone reproduction, potential nesting sites are abundant (e.g. fold in the kelp) and should not limit male reproduction, and finally males that already have a nest with eggs are still able to court females to increase the size of their current brood.

## 3. Behavioural analysis framework

### 3.1 General framework

Mating related behaviours require encounters between individuals. Therefore, we assume the number of occurrences *B* of a given behaviour to be the product of the number of encounters between the relevant individuals, *E*, with the probability for the behaviour to arise during an encounter, *P*, which itself is a function of the propensity of individuals to perform the behaviour, *Prop*.

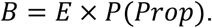

The relationship between the probability for the behaviour to arise and the propensity for individuals to behave will be clarified when specific examples are given in the next section (4.2). In the general case, we assume that the number of encounters between relevant individuals is proportional to the number of individuals present:

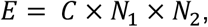

where *C* is a proportionality constant, and *N*_1_ and *N*_2_ are the respective numbers of individuals of the two classes. The product of individual numbers is replaced by *N*^2^ if there is only one class of individuals.

### 3.2 Framework adapted to the study

#### Courtship initiation behaviour

The number of female courtship initiations towards males, *B*_*fm*_, is the product of the number of encounters between a male and a female both ready to mate, *E*_*fm*_, times the propensity of females to initiate courtship when meeting a male, *Prop*_*fm*_.

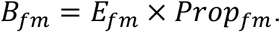

In the courtship case, the probability for the behaviour to arise and the propensity to perform the behaviour are the same: *P=Prop*. We will see that it is not the case for same-sex behaviours.

In practice, observations are made along transects of varying length and it is more convenient to deal with behaviour rates (behaviour per meter of transect), and with densities (fish per meter of transect) rather than numbers of individuals. With a transect of length *L*, we have:

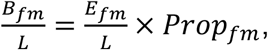

meaning that the rate of female courtship initiation is the product of male-female encounter rate and female propensity to court. Replacing numbers of individuals *N* by densities 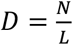,we obtain:

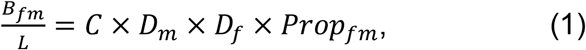

Where *C* represents an unknown proportionality constant. The same principle applies to male courtship.

#### Same-sex agonistic behaviour

The number of male-male agonistic behaviours, *B*_*mm*_, is the product of the number of encounters between two males, *E*_*mm*_, by the probability to observe at least one of the two males perform the behaviour. The probability for one male to perform the behavior being *Prop*_*mm*_, the probability that none of the two males perform a behavior is: (1 − *Prop*_*mm*_)^2^. Thus, in this case, the relationship between the probability for the behaviour to arise during an encounter and the propensity of a single individual to perform the behaviour is: *P*_*mm*_ = 1 − (1 − *Prop*_*mm*_)^2^. Then:

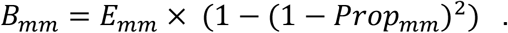

Following the same reasoning as for courtship, we transition to behaviour rates and obtain:

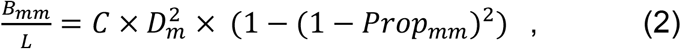

where *L* is the length of the observation transect and *C* is an unknown proportionality constant in the relationship between fish densities and encounter rates. The same principle applies to female same-sex agonistic behaviours.

### 3.3 Estimating propensities from field data

Equations (1) and (2) allow us to estimate behavioural propensities when the number of behaviour occurrences and individual densities are known. For example, with female courtship we can estimate propensity as:

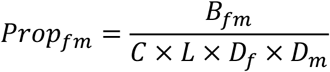

We note that the denominator: *C* × *L* × *D*_*m*_ × *D*_*f*_, is an estimate of the total number of encounters between males and females, which we later use as a weight variable in our binomial model (see Methods 4.4). Applying the same logic for male agonistic behaviours we obtain:

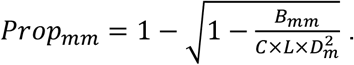

This solution is valid as long as the number of behaviour occurrences *B*_*mm*_ is not superior to the estimated total number of encounters: 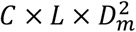, which should always be true. *C* is an unknown proportionality constant, defining the relationship between individual densities and encounter rates.

#### Choice of the proportionality constant

For convenience, when analysing the field data, we set one value for *C* for each type of behaviour (courtship or agonistic) so that the highest propensity in each dataset is 1. This way, propensities are scaled and can be analysed as a binomial variable (see Methods 5.4). This means that (i) propensities are not comparable across behaviour types, but remain comparable across sexes and time periods within each behaviour type, and (ii) we present scaled propensities, where the maximum propensity in the data is equal to one, whereas real propensities in the field may be lower (the true maximum propensity is not known).

### 3.4 Social environment proxy and behavioural reaction norms

We assume that individuals perceive their social environment and adjust their behaviour propensity following a reaction norm. The OSR is a classic proxy for the social environment used to predict mating competition and associated behaviour. The OSR is usually calculated as the relative number of males and females in the mating pool (Kvarnemo and Ahnesjö 1996, Box 1), but can also be calculated as a relative proportion of male to female densities:

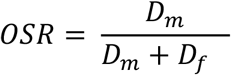

This formulation is insightful, because male and female densities are proportional to male and female encounter rates, from the perspective of a focal fish. Consequently, OSR can be understood as a comparison of the encounter rates of potential mates and competitors from an individual fish perspective, making it a plausible cue for reaction norms of sexual display behaviour. We describe these reaction norms as the relationship between the propensities calculated in 4.2 and the OSR, where the trait is the propensity to behave and the environment is the OSR (see Methods 5.4 for the statistical method).

#### Behavioural switching point

calculated as a proportion, the OSR is bound between 0 and 1, and when OSR<0.5, females are more abundant than males in the mating pool. However, depending on the mating system, individuals may not exit the mating pool after one mating only (collateral investment, Parker and Simmons 1996). For example, in our study species it was estimated that males in the Kristineberg population can on average hold the egg clutches of 4-5 females in their nest at a time (Mobley et al. 2009, Monroe et al. 2016). This can of course vary across populations, but regardless of the exact number, an OSR of 0.5 may not be perceived as balanced by the individual fish if they are able to integrate the collateral investment in their assessment of the social environment. With a remating coefficient *Rm*=4, the behaviour switching point could be when *D*_*f*_ = 4 × *D*_*m*_, which corresponds to OSR=0.2.

We illustrate the two potential switching points in the figures of the manuscript, OSR=0.5 and OSR=0.2. They can be seen as two limit cases, from the fish completely ignoring the collateral investment (OSR=0.5), to the fish acknowledging it fully (OSR=0.2), whereas the true perception of the social environment can be anywhere in between. We note that experimental evidence from the sand goby *P. minutus* shows that, in that closely related species, the collateral investment does not affect mating competition (Nyman et al. 2006).

## 4. Statistical Analysis

### 4.1 Behaviour rates

The behavioural rate corrected for density is, for each sex, the number of times a behaviour is observed per meter of transect, divided by the number of fish of the appropriate sex per meter of the same transect. The effects of time period, population and sex on the behaviour rates corrected for density were analysed using a generalised linear model with binomial response in *R* (R Core Team 2023) with the *glm* function. The binomial response was coded with the two outcomes (“failure” and “success” in the classical binomial framework) being the count of a specific behaviour, and the count of individuals minus the count of behaviours. The logic is that the count of individuals represents the number of encounters, and the count of behaviours the number of successes, the difference of the two the number of failures. Two separate models were fitted, one for courtship initiation and one for same-sex agonistic behaviours. Fixed effects in the models were time period, population and sex, with the three two-way interactions and the three-way interaction added. Post-hoc tests were executed with the *emmeans* function of the *emmeans* package (Russell 2023) in *R*. In this part of the analysis, time period was coded as a continuous variable, in order to model the temporal trajectories of behaviour rates.

### 4.2 Social environment

The effect of time period and population on the social environment proxy OSR was analysed with a generalised mixed effect model, using the *lme4* package (Bates et al. 2015) in R. The social environment proxy OSR was described as a binomial response variable, with the number of males and females observed in the mating pool representing the two outcomes. Time period (early, mid-, or late season) and population were fixed effects in the model, while the identity of sampling sublocation was fitted as a random effect. Pairwise comparisons of the social environment between each possible pair of population and time period were made using the *emmeans* package in R, with the Tukey method to adjust p-values.

### 4.3 Adult sex-ratio

The effect of time period and population on the proportion of males in the adult population (not restricted to the mating pool) was analysed with a generalised mixed model, using the *lme4* package in R. The response variable was coded as binomial with the number of adult males and the number of adult females representing the two outcomes. Time period (early, mid-, or late season) and population were fixed effects in the model, while the sampling sublocation was fitted as a random effect. Post-hoc tests were done with the *lstrends* function of the *lsmeans* package (Russell 2016) and *emmeans* in *R*.

### 4.4 Reaction norms

The behavioural reaction norms were estimated with a logistic regression of the propensity to perform a behaviour on the social environment proxy OSR. Two separate models were fitted, one for propensity to initiate courtship and one for propensity to same-sex agonistic behaviour. To model these, we fitted a binomial generalised mixed effect model, using the *lme4* package in *R*. The propensity to perform a behaviour was the response variable, weighted by the estimated number of encounters between relevant individuals (see Methods 3.3). The social environment (OSR), sex of the courting individual, and time period were fitted as fixed effects, with the addition of the OSR by sex and sex by time period two-way interactions. We note that the effect of time period was *a priori* not to be included in the model, since temporal changes in reaction norms were not expected. However, after visual exploration of the data, it was decided to include the effect of time period in the model. The identity of sampling sublocation was fitted as a random effect. In the case of courtship propensity, model comparison suggested to include the three-way interaction in the model, although statistically not significant, whereas it was excluded for agonistic propensity (Supplementary tables 1 and 2 for model comparisons). In the case of agonistic behaviours, one outlier was removed from the female data prior to the analyses because it had very low individual density. For both types of behaviour, five outlier points were removed either because they represented outliers in fish density (4/5), or in extreme OSR value (1/5), leading to improvement of the residual distribution (Supplementary tables 1 and 2). The data points excluded for the analysis are visible on Figures 5 and 6, and results are presented both with and without outliers included.

## Results

### Rates of sexual display behaviour over time

#### Courtship behaviours

The rate of courtship initiation by males and females varied over time in each population (Figure 2). Modelling the effect of population, time period and sex on the rate of courtship initiation, we found a significant three-way interaction between these effects (Table 1). Post hoc tests showed that in Austevoll, females had an overall higher courtship initiation rate than males (pairwise contrast: Estimate=1.65, SE=0.68, p=0.016), whereas in Hitra females had a lower courtship initiation rate than males (pairwise contrast: Estimate=-2.75, SE=0.44, p<0.001). Temporal trajectories by population showed that male courtship initiation rate decreased over time in all populations, while females showed a significant increase over time in the two southernmost populations of Kristineberg and Arendal, a significant decrease in the two intermediate populations of Austevoll and Hitra, and no courtship initiation at all in the two northernmost populations (Helligvær and Ringstad, Figure 2, Table 2).

**Figure 2.**
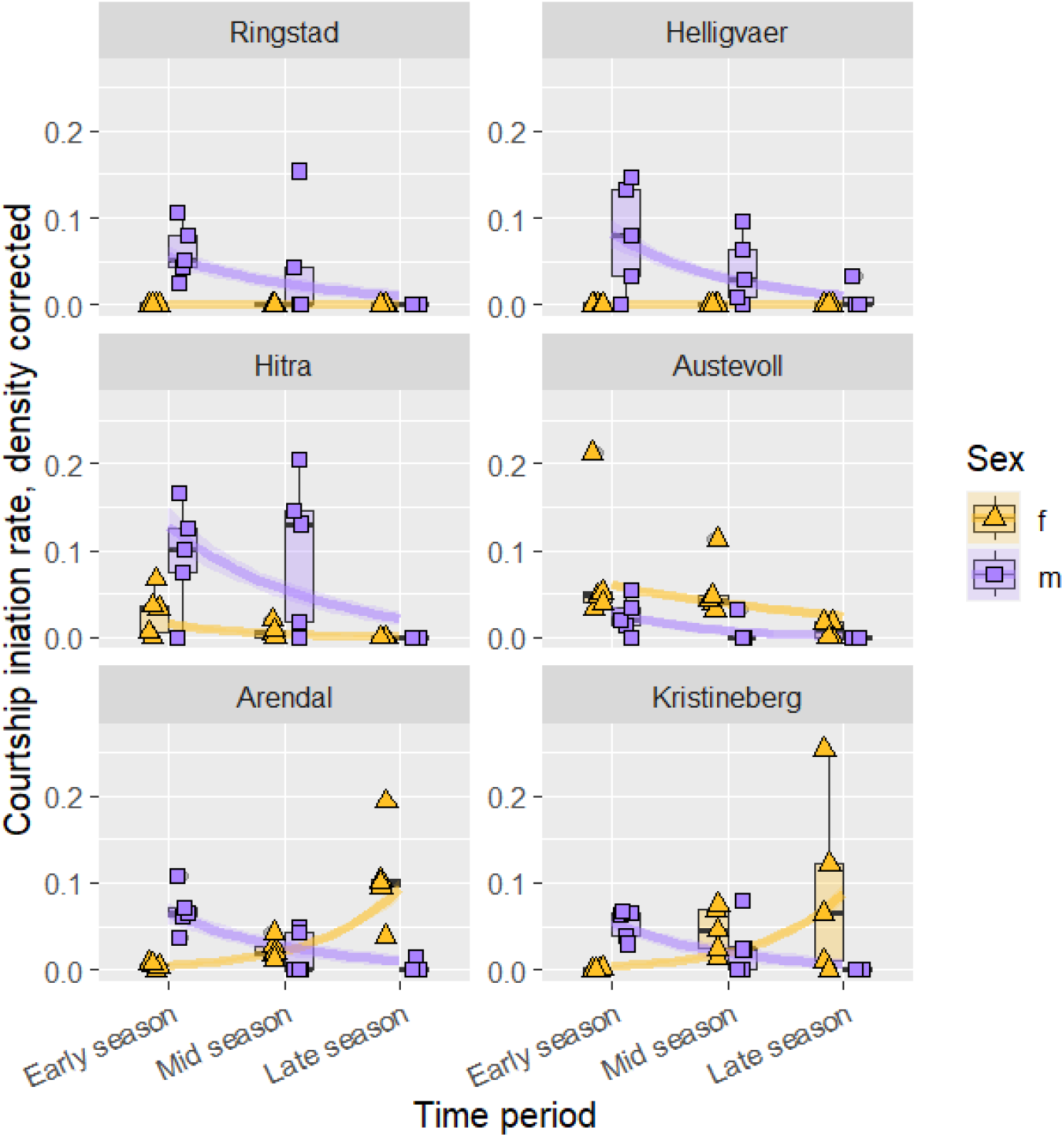
Density corrected rate of courtship initiation by either females (gold) or males (purple) for each population of two-spotted goby (Pomatoschistus flavescens) and time period. The panels are arranged from North to South in reading order. Each dot represents one sampling occurrence, with 5 sublocations for each population and time period. For each sex, the frequency of courtship initiation is measured per meter of transect, and divided by the local density of the respective sex (number of fish per meter of transect). Purple squares represent male rates, and gold triangles represent female rates. Boxes encompass 50% of the data around the median (dark bar), while whiskers indicate 1.5 times the interquartile range. Trend lines show the fit of the binomial generalised linear model, and shaded areas the standard error around the fit.

**Figure 3.**
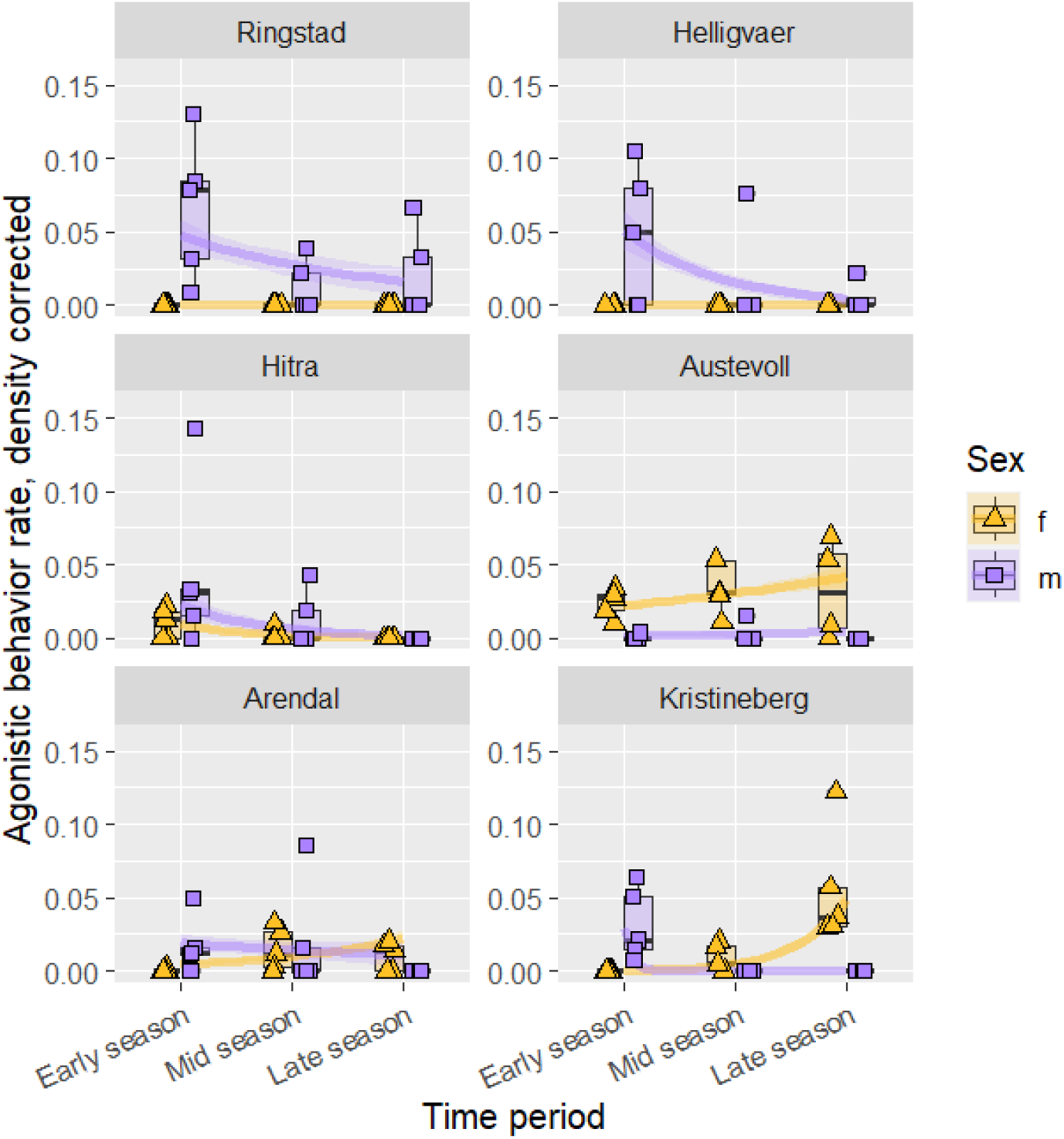
Density corrected rates of same-sex agonistic behaviours performed by either females (gold) or males (purple) for each population of two-spotted goby (Pomatoschistus flavescens) and time period. The panels are arranged from North to South in reading order. Each dot represents one sampling occurrence, with 5 sublocations for each population and time period. For each sex, the frequency of agonistic behavior initiation is measured per meter of transect, and divided by the local density of the respective sex (number of fish per meter of transect). Purple squares represent male rates, and gold triangles represent female rates. Boxes encompass 50% of the data around the median (dark bar), while whiskers indicate 1.5 times the interquartile range. Trend lines show the fit of the binomial generalised linear model, and shaded areas the standard error around the fit.

**Table 1:** Anova table (type 3) for the binomial generalized linear model of the effect of time period, population and sex on the rate of courtship initiation, corrected for density.

| Effect | Chi square | Df | P value |
| --- | --- | --- | --- |
| Time period | 122 | 1 | <0.001 |
| Sex | 72 | 1 | <0.001 |
| Population | 240 | 5 | <0.001 |
| Time period:Sex | 52 | 1 | <0.001 |
| Population:Sex | 26 | 5 | <0.001 |
| Time period:Population | 202 | 5 | <0.001 |
| Time period:Sex:Population | 33 | 5 | <0.001 |

**Table 2:** Slope estimates for male and female courtship initiations rate over time, for each population. Courtship rate is the response variable (binomial), population, sex and time period are fixed effects. As no courtship was observed at Helligvær and Ringstad, slope, SE and P-values are not applicable (N/A) for these locations.

| Population | Sex | Slope | SE | P value |
| --- | --- | --- | --- | --- |
| Kristineberg | F | 1.55 | 0.16 | <0.001 |
|  | M | -1.07 | 0.49 | 0.030 |
| Arendal | F | 1.52 | 0.17 | <0.001 |
|  | M | -0.95 | 0.36 | 0.009 |
| Austevoll | F | -0.46 | 0.12 | 0.001 |
|  | M | -1.39 | 0.72 | 0.052 |
| Hitra | F | -1.51 | 0.46 | 0.001 |
|  | M | -0.92 | 0.23 | <0.001 |
| Helligvær | F | N/A | N/A | N/A |
|  | M | -1.03 | 0.26 | <0.001 |
| Ringstad | F | N/A | N/A | N/A |
|  | M | -0.87 | 0.38 | 0.021 |

#### Same-sex agonistic behaviours

Modelling the effect of population, time period and sex on the number of agonistic events, with a binomial generalised linear model, we found a significant three-way interaction between these effects (Table 3). Post hoc tests showed that in Austevoll, females performed overall more agonistic behaviours than males (female-male pairwise contrast, Austevoll: Estimate=2.46, SE=0.89 p=0.006), while in the other populations where both sexes exhibited agonistic behaviours no significant sex-differences were found. In the two northernmost populations (Helligvær and Ringstad), female agonistic behaviours were not observed. Temporal trajectories by population showed a significant decrease of behaviour rate over time for males in Helligvær (Marginal mean of full model, Slope=-1.23, SE=0.37, p<0.001), a decrease for females in Hitra (Slope=-1.82, SE=0.70, p=0.011), and an increase over time for females in the three southernmost populations (Kristinberg: Slope=2.31, SE=0.36, p<001, Arendal: Slope=0.80, SE=0.25, p= 0.001, Austevoll: Slope=0.33, SE=1.13, p=0.01).

**Table 3:** Anova table (type 3) for the binomial generalized linear model of the effect of time period, population and sex on the count of agonistic events.

| Effect | Chi square | Df | P value |
| --- | --- | --- | --- |
| Time period | 87.1 | 1 | <0.001 |
| Sex | 64.8 | 1 | <0.001 |
| Population | 81.7 | 5 | <0.001 |
| Time period:Sex | 51.4 | 1 | <0.001 |
| Population:Sex | 35.0 | 5 | <0.001 |
| Time period:Population | 66.4 | 5 | <0.001 |
| Time period:Sex:Population | 22.1 | 5 | <0.001 |

### Social environment (OSR) over time

We expect the OSR to affect encounter rates, as well as the propensity of each sex to perform behaviours due to their own perception of the relative encounter rates of potential mates and competitors (see Methods 4.4). The OSR is calculated as the relative proportion of the densities of males and females ready to mate, which are plotted separately in Supplementary Figure S2. Here we show how OSR varied over time in each population (Figure 4).

**Figure 4.**
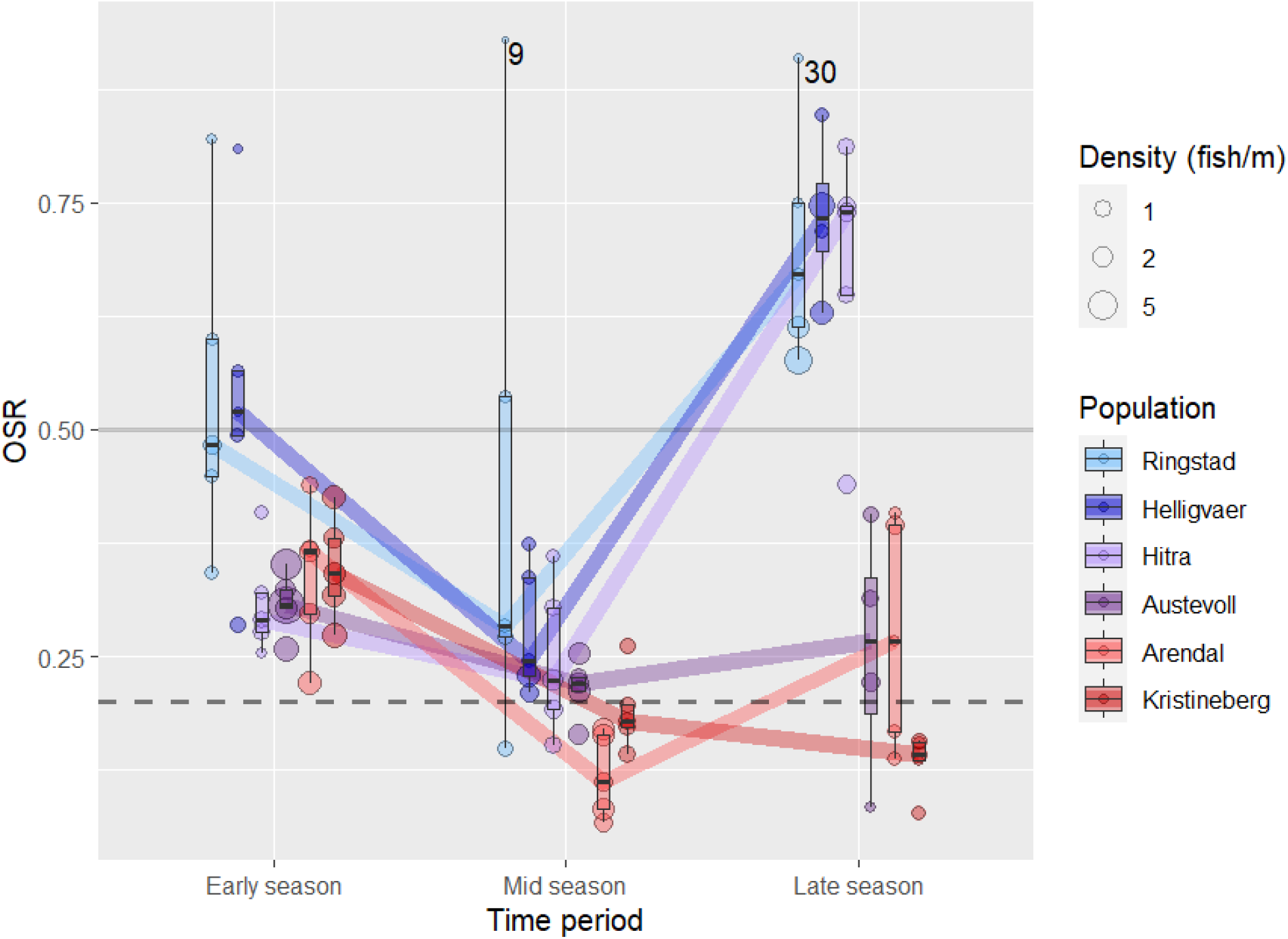
Temporal trajectories of the operational sex ratio (OSR) in the different study populations. Each dot represents a population census event, with each study population replicated in 5 sublocations at each time period. The size of the dots represents population density (number of fish/meter), the colour refers to the population of origin. Trend lines are connecting the medians of each population across time. The solid grey horizontal line represents the switch point between males (OSR>0.5) or females (OSR<0.5) being more abundant in the mating pool. The dashed horizontal line represents the expected behavioural switching point with a when accounting for collateral investment (see Methods 4.4). For transects where 30 fish or less were observed in total, the number of fish is indicated next to the dot. Supplementary Figure S4 shows the trendline for each sub-location instead of at the population level.

**Figure 5.**
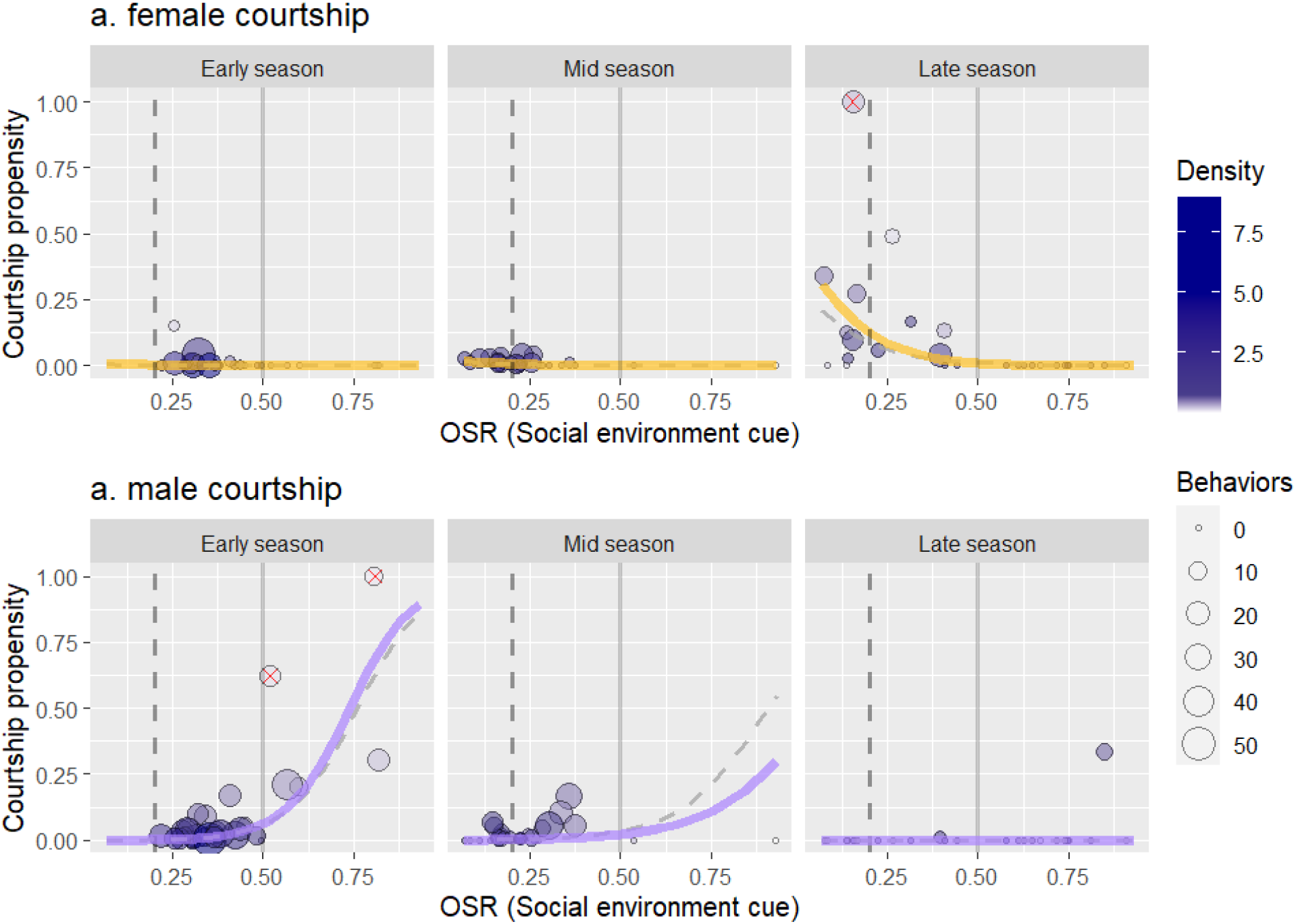
Sex-specific reaction norms for the propensity to initiate courtship as a function of the operational sex ratio (OSR), for each time period. Each dot is a sublocation at a given sampling time (representing two consecutive observation transects), the size of the dots corresponds to the number of courtship initiation behaviours observed for the focal sex on that transect. The colour fill of the dots reflects the local density (individuals per meter of transect), where the white fill indicates low density (>0,75 fish/m) leading to higher stochasticity in the estimation of propensity. Courtship propensity is the proportion of male-female encounters leading to courtship initiation by the focal sex (see Methods 4.3). The solid grey vertical line represents the switch point between males (OSR>0.5) or females (OSR<0.5) being more abundant in the mating pool. The dashed vertical line represents the switching point between males or females being perceived as more abundant in the mating pool, taking into account the collateral investment (see Methods 4.4). Solid coloured trend lines represent predictions from a binomial generalised mixed-model fitted to the data. Data points marked with a red cross (5 in total) represent potential outliers, and the grey dashed trend lines represent model predictions when the outliers are excluded.

**Figure 6.**
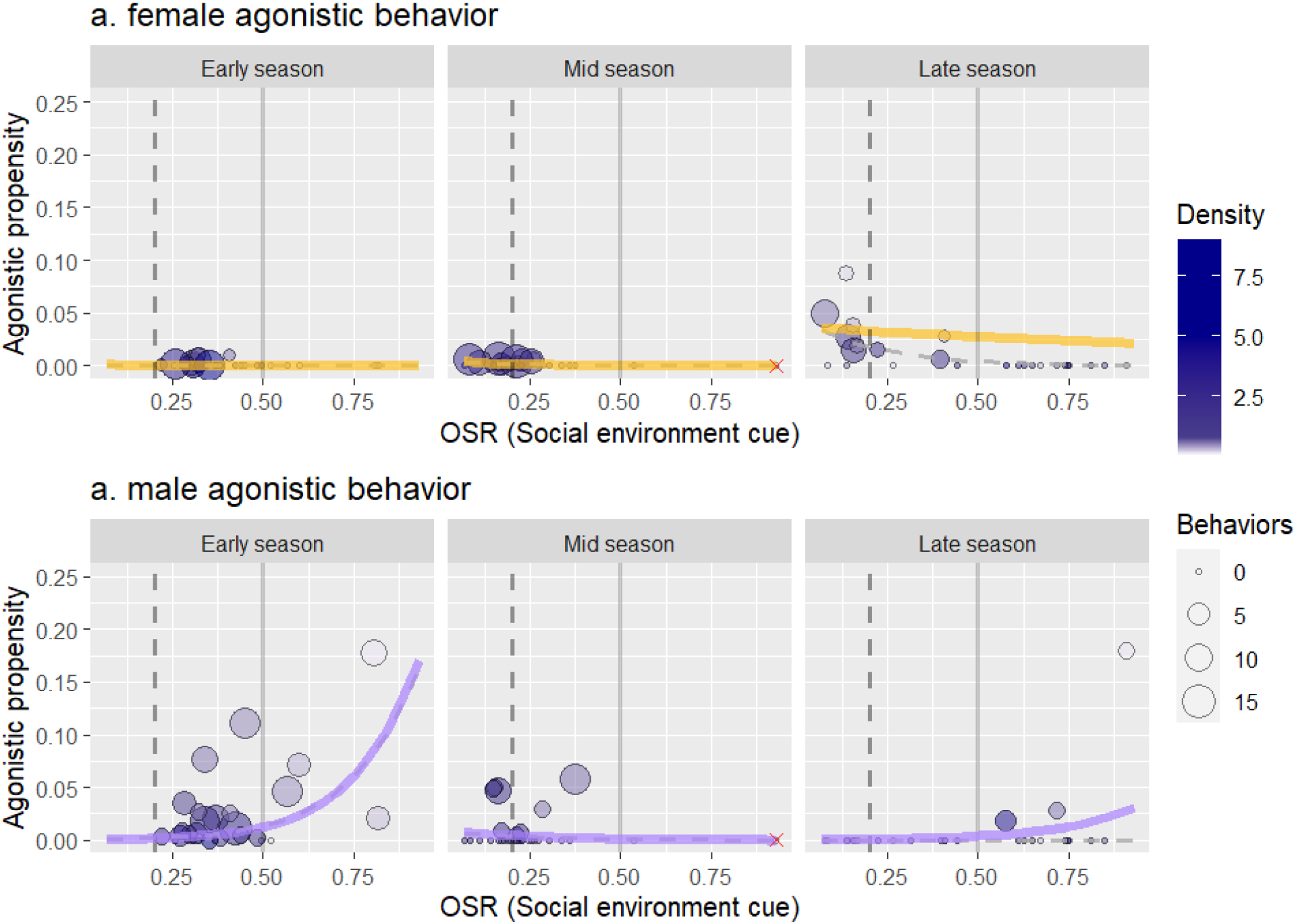
Sex-specific reaction norms for the propensity to perform same-sex agonistic behaviour as a function of operational sex ratio (OSR), for each time period. Each dot is a sublocation at a given sampling time (representing two consecutive observation transects), the size of the dots corresponds to the number of agonistic behaviours observed for the focal sex on that transect. The colour fill of the dots reflects the local density (individuals per meter of transect), where the white fill indicates low density (>0,75 fish/m) leading to higher stochasticity in the estimation of propensity. Agonistic propensity is estimated from the proportion of same-sex encounters leading to agonistic behaviour for the focal sex (see Methods 4.3). The solid grey vertical line represents the switching point between males (OSR>0.5) or females (OSR<0.5) being more abundant in the mating pool. The dashed vertical line represents the switching point between males or females being perceived as more abundant in the mating pool, taking into account the collateral investment (see Methods 4.4). Solid coloured trend lines represent predictions from a binomial generalised mixed-model fitted to the data. Data points marked with a red cross (5 in total) represent potential outliers, and the grey dashed trend lines represent model predictions when the outliers are excluded. For better visualisation, the y-axis was restricted to the 0-0.25 range, leading to the exclusion of 3 outlier points. The full range is visible in Supplementary Figure S7.

Time period, population, and their interaction all affected the social environment (Table 4). In particular, all populations presented a drop in OSR mid-season (Figure 4), indicating the presence of numerous females in the mating pool (OSR is lower when females are more abundant). A post-hoc test confirmed that mid-season OSR was significantly lower than both early and late season OSR (Table 5).

**Table 4:**
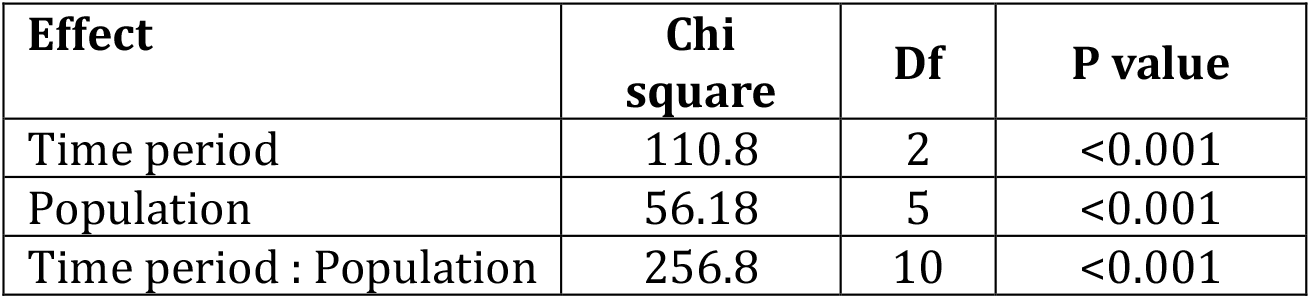
Type 3 Anova table for the mixed effect generalized linear model of the effect of population and time period on social environment (OSR), with sampling location as a random effect.

| Effect | Chi square | Df | P value |
| --- | --- | --- | --- |
| Time period | 110.8 | 2 | <0.001 |
| Population | 56.18 | 5 | <0.001 |
| Time period : Population | 256.8 | 10 | <0.001 |

**Table 5:** Pairwise comparison of social environment (OSR) across time periods, with Tukey’s adjustment for multiple testing.

| Comparison | Estimate | SE | Z ratio | P value |
| --- | --- | --- | --- | --- |
| Early-Mid | 0.86 | 0.045 | 19.0 | <0.0001 |
| Early-Late | -0.24 | 0.066 | -3.63 | 0.0008 |
| Mid-Late | -1.10 | 0.068 | -16.1 | <0.0001 |

OSR fluctuations are the outcome of two different processes: changes in ASR, and changes in the egg maturation stage of females over time. ASR became increasingly female-biased in all populations over time, which may be due to higher male mortality (Supplementary Figure S5, Table 6). This relative decrease in the abundance of adult males contributed to the mid-season dip in OSR. However, the main driver of the mid-season dip was likely the peak in female egg maturity at that time period (data shown in Martinossi-Allibert et al, unpublished data).

**Table 6:** Slope estimates for adult sex ratio over time from early to late season, for each population. The proportion of males is the response variable (binomial), population and time period are fixed effects and sampling sublocation is fitted as a random effect.

| Population | Slope | SE | Z ratio | P value |
| --- | --- | --- | --- | --- |
| Kristineberg | -0.46 | 0.063 | -7.25 | <0.001 |
| Arendal | -0.51 | 0.065 | -7.83 | <0.001 |
| Austevoll | -0.62 | 0.062 | -10.0 | <0.001 |
| Hitra | -0.24 | 0.058 | -4.20 | <0.001 |
| Helligvaer | -0.88 | 0.056 | -15.73 | <0.001 |
| Ringstad | -1.12 | 0.061 | -18.5 | <0.001 |

The two northernmost populations (Helligvær, Ringstad) generally showed more male-biased mating pools. In the last time period, the three populations to the north presented significantly fewer females in the mating pool than the three populations to the south, which indicates that female reproduction stopped earlier in the north. The p-values of all pairwise comparisons between each population by time period combination are shown in Supplementary Figure S6.

### Sex-specific behavioural reaction norms

#### Courtship behaviours

The estimated propensity for each sex to initiate courtship for each sampling occurrence allowed us to model behavioural reaction norms with respect to OSR for each sex and time period (Figure 5). We modelled the propensity to initiate courtship as a binomial response to OSR, sex and time period and relevant interaction effects, with the addition of sampling sublocation as a random effect (Table 7). This model confirmed opposite slopes for the two sexes, with male propensity increasing with OSR and female propensity increasing with decreasing OSR (marginal means estimated from the full model for females: slope=-7.44, SE=0.75, and males: slope=11.1, SE= 0.99). In addition, males had higher propensity to initiate courtship early and mid season, but females had higher propensity in late season (Contrasts female - male: Early=-1.74, Mid=-3.21, Late=3.52, p values for all three contrasts <0.05, after Tukey adjustment). The three-way interaction in the model is close to the significance threshold (Table 4, p=0.061), which gives mild support to the slopes of the reaction norms differing across sexes and timepoints, in addition to the intercepts (Figure 5). In mid-season, we note that the limited range of the social environment observed did not allow to draw the full reaction norm (OSR was highly female biased in most populations).

**Table 7:** Anova table (type 3) for the binomial generalized linear mixed effect model of the effects of social environment (OSR), time period and sex on propensity to initiate courtship, outliers excluded.

| Effect | Chi square | Df | P value |
| --- | --- | --- | --- |
| Intercept | 88.7 | 1 | <0.001 |
| OSR | 1.76 | 1 | 0.184 |
| Sex | 24.0 | 1 | <0.001 |
| Time period | 83.0 | 2 | <0.001 |
| Sex:Time period | 3.52 | 2 | 0.17 |
| OSR:Sex | 47.1 | 1 | <0.001 |
| OSR:Sex:Time period | 9.01 | 4 | 0.061 |

Together, this means that the courtship propensity of each sex generally responded to the social environment as predicted by OSR theory, with the courtship propensity of each sex increasing with its own abundance in the mating pool. Moreover, the sexes varied seasonally in their propensity to initiate courtship, with males having a high propensity to initiate courtship early in the season, while females did so only late in the season (Figure 5).

#### Same-sex agonistic behaviours

Overall, propensities to perform agonistic behaviours appeared lower than propensities to perform courtship (comparing Figure 5 and 6). Modelling the propensity to perform same-sex agonistic behaviour as a binomial response to OSR, sex and time period, with relevant interaction effects, and sampling sublocation as a random effect (Table 8), confirmed opposite slopes for the two sexes, with male propensity increasing with increasing OSR and female propensity increasing with decreasing OSR (marginal means estimated from the full model for females: slope=-3.99, SE=1.64, and males: slope=5.32, SE= 1.14). In addition, males had a higher propensity to perform agonistic behaviours in early and mid-season, while females had a higher propensity in late season (female - male contrasts: Early=-2.19, Mid=-2.95, Late=2.42, p values for all three contrasts <0.05, after Tukey p-value adjustment).

**Table 8:** Anova table (type 3) for the binomial generalized linear mixed effect model of the effects of social environment (OSR), time period and sex on propensity to perform same-sex agonistic behavior.

| **Effect** | **Chi square** | **Df** | **P value** |
| --- | --- | --- | --- |
| Intercept | 110 | 1 | <0.001 |
| OSR | 5.92 | 1 | 0.015 |
| Sex | 3.00 | 1 | 0.083 |
| Time period | 68.8 | 2 | <0.001 |
| Sex:Time period | 37.4 | 2 | <0.001 |
| OSR:Sex | 22.4 | 1 | <0.001 |

The three-way interaction between OSR, sex and time period was not significant and excluded from the final model. Therefore, in the case of agonistic behaviours, the sexes show differences in the intercept of the reaction norm over time (change in average propensity), but there is no evidence for a change in slope.

Together, this means that similarly to courtship behaviours, the propensity for each sex to perform agonistic behaviours responded to the social environment as predicted by OSR theory, with males and females showing overall higher propensity when their own sex was more abundant in the mating pool. The sexes varied seasonally in their propensity to perform agonistic behaviours, with males having higher mean propensity than females early and mid-season, but lower than females late in the season.

## Discussion

This work is, to our knowledge, the first study comparing temporal trajectories of sexual display behaviours across several populations experiencing different environmental conditions. We argue that such a spatio-temporal scale is needed to fully grasp the relationship between ecological variables, social environment (OSR) and mating competition, and indeed our data provides novel insights into these complex dynamics through the *P. flavescens* system. First, we confirmed the main prediction of OSR theory: the propensity to perform sexual display behaviours in the field increased for males and females with the relative presence of their own sex in the mating pool. We also discovered that the propensity to perform these behaviours was overall higher for males early in the breeding season, and higher for females late in the breeding season, irrespective of the OSR.

The temporal variation of sex-specific reaction norms significantly updates our understanding of the *P. flavescens* system, which is a prominent model system for OSR studies in the field. Earlier work by Forsgren et al. (2004) suggested, with data from only one population, sex-specific reaction norms constant over time. The population studied by Forsgren et al. (2004) was sampled again in the present study, leading to very similar observations. In order to reveal the temporal variation in reaction norm, it was necessary to observe a range of OSR at each sampling time, which was only made possible by studying several populations following different temporal trajectories of OSR.

Although all the populations conformed to the same behavioural reaction norms, they had very different temporal trajectories of sexual display behaviour throughout the breeding season. For example, females from the two northernmost populations were not observed performing a single courtship initiation or agonistic behaviour throughout the season, while they did so frequently elsewhere.

Such discrepancies remind us that we should not expect mating competition to be fixed within species or even within populations over time. Indeed, sexual selection theory states that the sexes are limited by different resources (e.g., Emlen and Oring 1977), and thus changes in ecological conditions should be expected to have sex-specific effects, tipping the balance of mating interactions (e.g., Zikovitz and Agrawal 2013, Martinossi-Allibert et al. 2020, Hare and Simmons 2020). In marine fish with nest-guarding males, akin to *P. flavescens*, the literature provides three striking examples where a shortage of nesting sites in natural populations leads to fewer males in the mating pool (female-biased OSR) and in turn to intensified female courtship and female-female competition (Almada et al. 1995, Borg et al. 2002, Garcia-Berro et al. 2019). These studies show how one particular environmental parameter can affect OSR and in turn sexual display behaviour. In the next paragraphs, we discuss the ecological and demographic variables that are specific to our study system and likely to drive OSR fluctuations, and in turn sexual display behaviour along the latitudinal gradient of the Norwegian coastline.

The breeding window for marine ectotherms, defined by water temperature and primary productivity, narrows down with increasing latitude along the Norwegian coastline (Martinossi-Allibert et al., unpublished data). As a result, spawning was still ongoing in southern populations in late season, while it had mostly stopped in northern ones (Martinossi-Allibert et al., in prep). This means that at the last time period when female propensity for sexual display behaviour is the highest, female-biased OSR are only found in southern populations. Thus, a latitudinal change in environmental conditions, which affects the timing of spawning, likely contributed to the complete absence of female sexual display behaviours in the two northernmost populations, and their decrease over time in the two intermediate populations. Among the very few studies that have explored latitudinal effects on mating competition, Monteiro and Lyons (2012) and Monteiro et al. (2017) have compared proxies of sexual selection and reproductive phenology of the worm pipefish *Nerophis lumbriciformis* across its range in the Northeast Atlantic. They showed that the breeding window for the species decreased with latitude, as for *P. flavescens*. However, their data indicated higher female competition at both the northern and southern ends of the species distribution, while we saw no female competition in the North, close to the range limit. Both *N. lumbriciformis* and *P. flavescens* have paternal care of eggs, but the pouch-brooding of male pipefish imposes a hard limit on male potential reproductive rate in that species, which does not exist in *P. flavescens*. Fujimoto et al. (2015), although not describing temporal trajectories, compared sexual display behaviours of the medaka fish (*Oryzias latipes*, no parental care) from different latitudes. They found that lower latitude males were more eager to compete and court, which they attributed to a generally male-biased OSR, due to a longer breeding period at low latitude. It is striking to note that in these two cases of latitudinal gradients, as well as in the present study, the narrowing down of the breeding window with latitude comes into play. However, the three species have different mating systems, in particular different modes of parental care, and the resulting effects on OSR and mating competition are truly at odds.

Adult Sex Ratio (ASR) biases contribute to OSR biases. From earlier studies in the southern population of Kristineberg, the ASR of *P. flavescens* in the breeding habitat is known to be largely female-biased (∼25% males on average), a bias that increases throughout the breeding period (e.g. Forsgren et al. 2004). This ASR bias has been attributed to higher male mortality prior to and during the breeding season (Forsgren et al. 2004), and a bias sex-ratio at birth is a possibility. However, in the two northernmost populations of our study, we observed an ASR close to balanced at the earliest time period, and recent genomic data suggests a purely genetic sex-determination system (Martinossi-Allibert et al. unpublished data). These new facts shed a different light on the female-biased ASR of southern populations. We propose that ASR in the breeding habitat at a given time may not be representative of the entire adult population, but that males could instead enter the breeding habitat sequentially over a period that varies depending on the length of the breeding window, itself determined by latitude. In northern populations, where breeding is concentrated in time, most males may enter the breeding habitat at once, leading to a balanced ASR. Whatever the underlying reason, the greater relative abundance of males in Northern populations could also contribute to the absence of female competition in these populations.

Male presence in the mating pool is difficult to assess in the field and we have chosen to include all adult males with clear secondary sexual characters in the mating pool. However, it is possible that only nest-holding males are qualified to mate, the proportion of which could be affected by male density in the breeding habitat (as suggested in Garcia-Berro et al. 2019 for the sand goby *P. minutus*). Fish density in general decreased with latitude, as well as nest occupancy of artificial nests placed in the field (Martinossi-Allibert et al., in prep). Higher nest availability in the North could have contributed as well to the absence of courtship by females, because of a higher proportion of adult males being nest-holders and thus effectively qualified to mate. In that case, male presence in the mating pool in southern populations could have been overestimated.

We propose one last ecological variable susceptible to affect sexual display behavior and varying with latitude. Garcia-Berro et al. (2019) suggest that low food abundance could also reduce female competition through effects on egg maturation in the close relative *P. minutus*. If that is the case *P. flavescens* as well, northern populations would again be expected to see reduced female competition as the female body condition factor decreased with latitude (Fulton’s condition factor, Martinossi-Allibert et al., in prep), likely reflecting lower food availability or an increased period of food shortage in the North.

The two southernmost populations showed the same pattern of early-season male courtship and late-season female courtship that had been observed in one of them in a study made about 20 years earlier (Forsgren et al. 2004). This suggests that although sexual display behaviour is plastic, it can also show consistency over time. This raises the question of whether this consistency in seasonal behavior trajectory across almost 20 years reflects stable environment conditions, causing the same plastic response, or if local adaptation could also play a role in this temporal stability. There are no genomics resources published in P. flavescens, but preliminary investigations suggest a connectivity break between the Skagerrak populations (Kristineberg, Arendal) and the northern ones (Martinossi-Allibert et al, unpublished data), which coincides with the end of late season female courtship. At the same time, environmental variables could explain the change in behaviour well, through effect on OSR, and common garden experiments would be required to address the issue fully.

In conclusion, we have shown that to grasp the complex interactions between ecological variables, social environment, and mating competition, it is necessary to compare populations across space and time. The two-spotted goby responded to its social environment by increasing courtship and agonistic behaviours when its own sex became more prevalent in the mating pool, which supports the main predictions of OSR theory. We were able to detect these patterns by accounting for the effect of male and female density on encounter rates, which varied widely throughout the breeding season and across latitudes.

*P. flavescens* is also known for performing ART, and such strategies are equally likely to change with latitude and associated environmental conditions. We suggest that a next step to further our understanding of this model system could be the quantification of sneaking events in the different populations. Our study also revealed that, regardless of the social environment, females had a higher propensity for sexual display behaviours late in the season, while males did so early on. Why this is the case could have to do with the scheduling of sex-specific reproductive tasks (maturing eggs for females, *versus* next guarding for males), or sex differences in prospects for winter survival of adults. Regardless, this finding calls for extreme caution when collecting data on sexual selection or reproductive behavior in the field, with respect to the timing of sampling.

