## Supplementary File F1 for "Sexual display behaviour follows consistent sex-specific reaction norms across latitude in response to operational sex-ratio"

### Supplementary File F1: The six study populations, 5 sub-locations in each, and sampling schedule.

**a. The 6 populations**

**b. The 5 sublocations for each population**

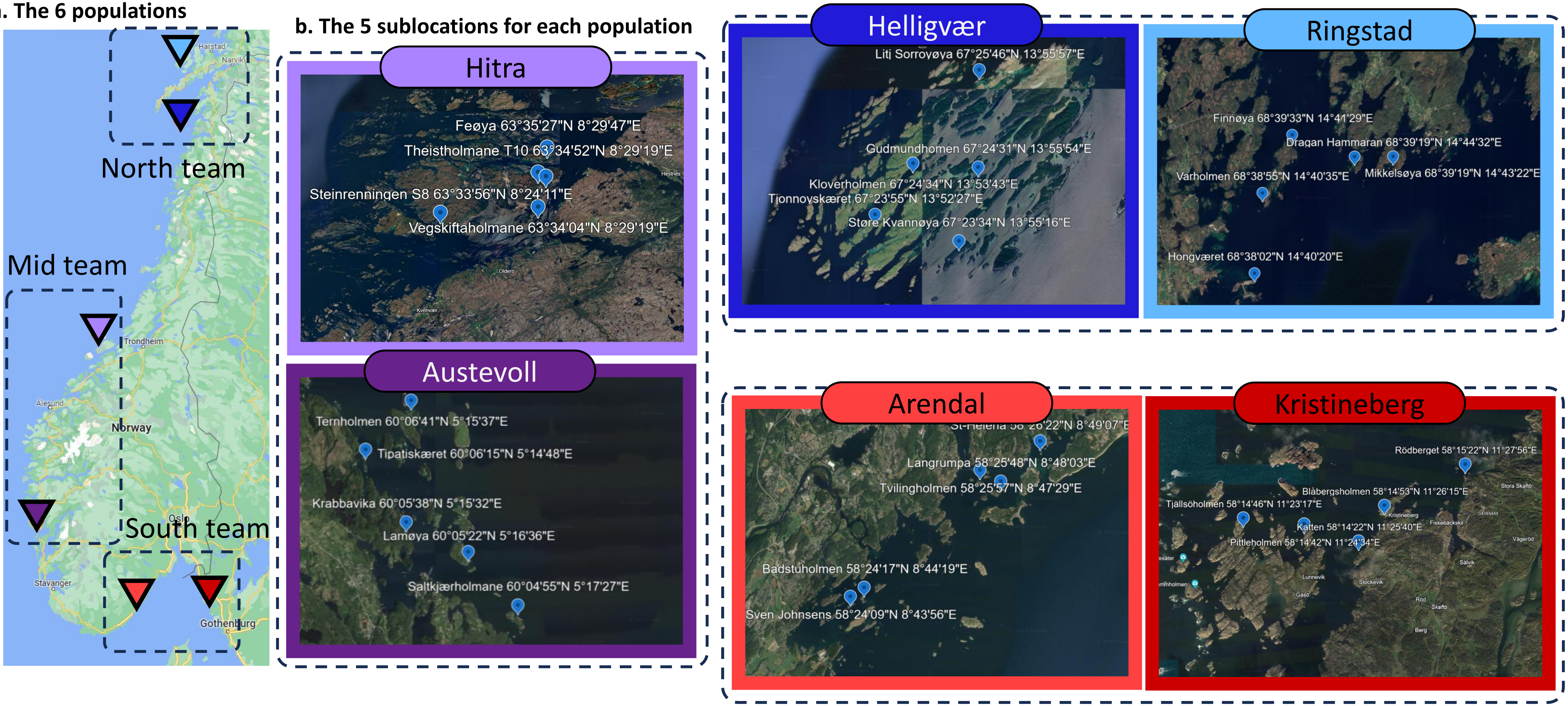


Maps from Google Earth 2023

**c. Timeline of the behavior and population census sampling for each sublocation**

Initials match the sublocation names in (b). Sampling was started later in northern locations to account for the expected delay in reproductive activity, matching delayed water temperature rise.

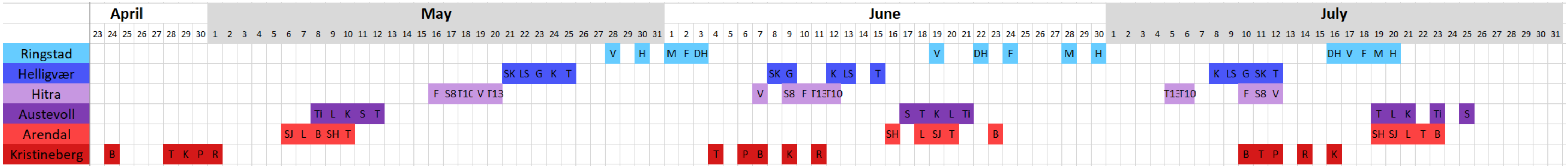

#### Table F1: Overview of transect length and GPS coordinate for the 30 sublocations

| Population | Sublocation | Transect length (m) | GPS.start.lat | GPS.start.long | GPS.end.lat | GPS.end.long |
|---|---|---|---|---|---|---|
| Kristineberg | Rodberget | 110 | 58N 15' 24'' | 11E 27' 60'' | 58N 15' 21.9'' | 11E 27' 58.2'' |
| Kristineberg | Katten | 110 | 58N14' 22.8'' | 11E 25' 39.3'' | 58N 14' 24'' | 11E 25' 36'' |
| Kristineberg | Blabergsholmen | 80 | 58N 14' 53'' | 11E 26' 15'' | 58N 14' 55'' | 11E 26' 11'' |
| Kristineberg | Pittleholmen | 125 | 58N 14' 39'' | 11E 24' 31'' | 58N 14' 43'' | 11E 24' 35'' |
| Kristineberg | Tjallsoholmen | 145 | 58N 14' 51'' | 11E 23' 18'' | 58N 14' 47'' | 11E 23' 18'' |
| Arendal | St Helena | 120 | 58N 26' 22'' | 8E 49' 07'' | 58N 26' 22'' | 8E 49' 09'' |
| Arendal | Langrumpa | 90 | N58.429912 | E8.8004622 | N58.4304093 | E8.8014198 |
| Arendal | Sven Johnsen | 105 | N58.4026569 | E8.7323643 | N58.4028706 | E8.7311090 |
| Arendal | Badstuholmen | 115 | 58N 24' 17'' | 8E 44' 19'' | 58N 24' 15'' | 8E 44' 22'' |
| Arendal | Tvillingholmen | 100 | N58.4326551 | E8.7915476 | N58.4328199 | E8.7902370 |
| Austevoll | Lamoya | 90 | N60 5' 23'' | E5 16' 36'' | N60 5' 22'' | E5 16' 33'' |
| Austevoll | Krabbavika | 55 | 60°05'38"N | 5°15'32"E | 60°05'39"N | 5°15'34"E |
| Austevoll | Saltkjaerholmane | 95 | N60 4' 56'' | E5 17' 26'' | N60 4' 55'' | E5 17' 23'' |
| Austevoll | Ternholmen | 80 | N60 06' 41'' | E5 15' 36'' | N60 06' 41'' | E5 15' 38.1'' |
| Austevoll | Tipatiskaeret | 80 | 60N 06' 16'' | 5E 14' 48'' | N60 06' 15.7'' | E5 14' 48'' |
| Hitra | Feoya | 115 | 63.5910059 N | 8.4966075 E | 63.5913066 N | 8.4987291 E |
| Hitra | Steinrenningen S8 | 90 | 63.5655958 N | 8.4031848 E | 63.5661535 N | 8.4036220 E |
| Hitra | Teistholmane T10 | 95 | 63.5813466 N | 8.4887322 E | 63.5815044 N | 8.4902962 E |
| Hitra | Vegskiftaholmane | 80 | 63.5678001N | 8.4887788 E | 63.5683827 N | 8.4889702 E |
| Hitra | Teistholmane T13 | 100 | 63°34'46"N | 8°29'39"E | 63.579837N | 8.4956878E |
| Helligvaer | Store-kvannoya | 210 | N67.393956 | E13.924549 | N67.392986 | E13.921224 |
| Helligvaer | Litj-sorroy | 150 | N67.42977547 | E13.93238484 | N67.43047742 | E13.93483948 |
| Helligvaer | Gudmunholmen | 240 | N67.408847 | E13.931860 | N67.407448 | E13.930830 |
| Helligvaer | Kloverholmen | 150 | N67.409678 | E13.895452 | N67.41049181 | E13.89812350 |
| Helligvaer | Tjonnoyskaeret | 155 | N67.398691 | E13.874389 | N67.398691 | E13.874389 |
| Ringstad | Varholmen | 115 | N68.6490484 | E14.6791256 | N68.648711 | E14.676340 |
| Ringstad | Hongvaeret | 110 | N68.633743 | E14.66941953 | N68.63395861 | E14.67244523 |
| Ringstad | Mikkelsoya | 225 | N68.65636073 | E14.72305055 | N68.65540936 | E14.72303402 |
| Ringstad | Finnoya | 140 | 68N 39' 31'' | 14E 41' 35'' | N68.65930773 | E14.69083428 |
| Ringstad | Dragan Hammaran | 130 | N68.656558 | E14.743009 | N68.655446 | E14.742129 |

**Transect coordinates for the 5 sub-locations of the Kristineberg population.**

Behavior and population census were recorded on transects indicated in yellow. Approximate transect length estimated from satellite image is given. Red lines indicate artificial nests lines.

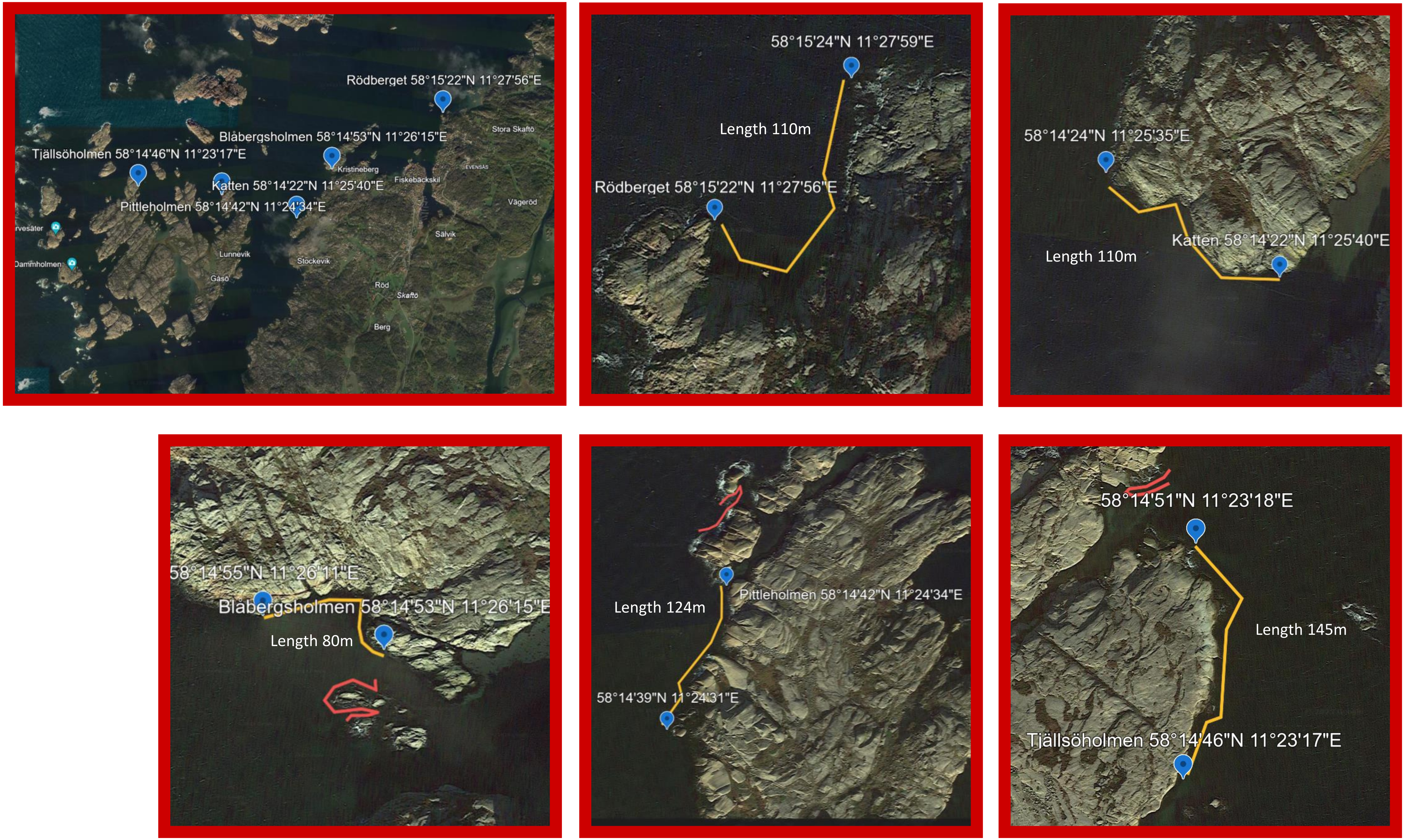


Images from Google Earth 2023

### Transect coordinates for the 5 sub-locations of the Arendal population.

Behavior and population census were recorded on transects indicated in yellow. Approximate transect length estimated from satellite image is given. Red lines indicate artificial nests lines.

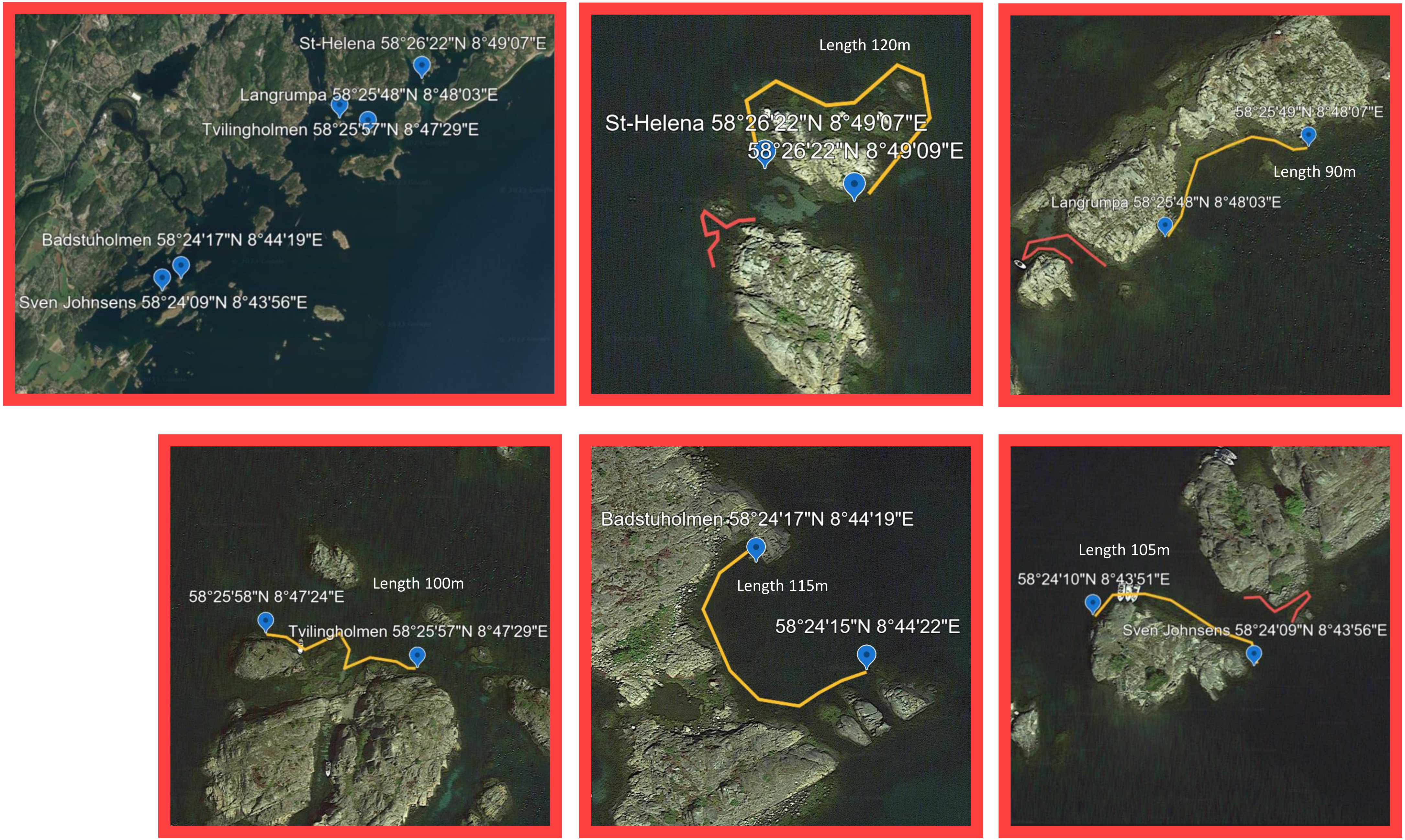


Images from Google Earth 2023

### Transect coordinates for the 5 sub-locations of the Austevoll population.

Behavior and population census were recorded on transects indicated in yellow. Approximate transect length estimated from satellite image is given. Red lines indicate artificial nests lines.

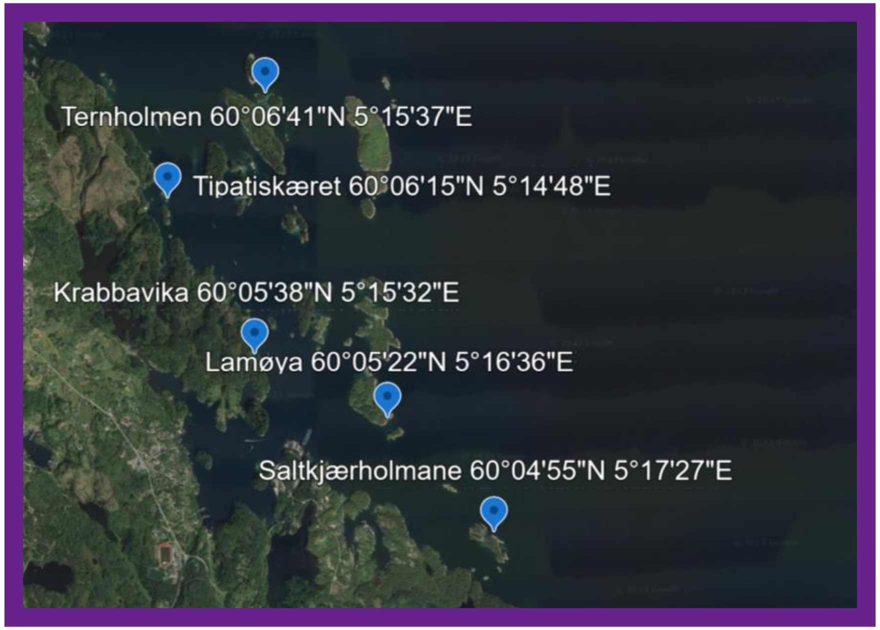


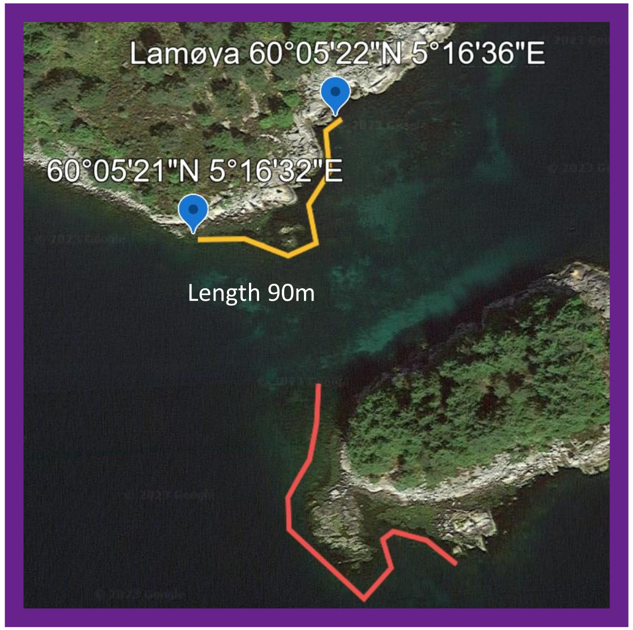


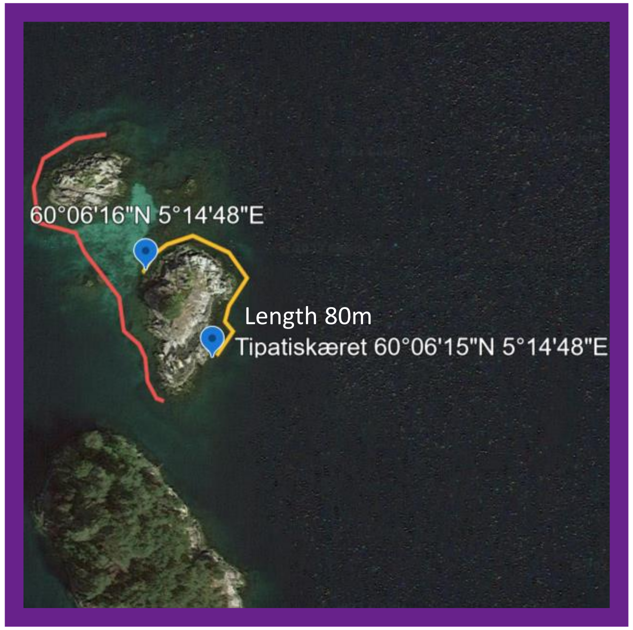


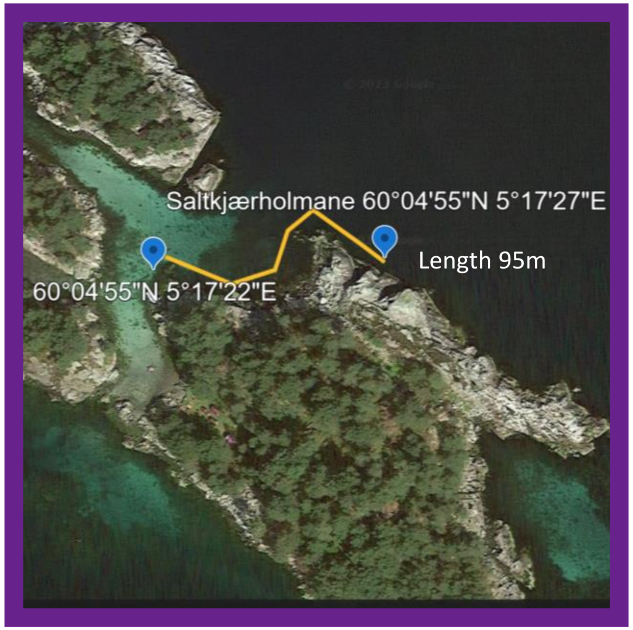


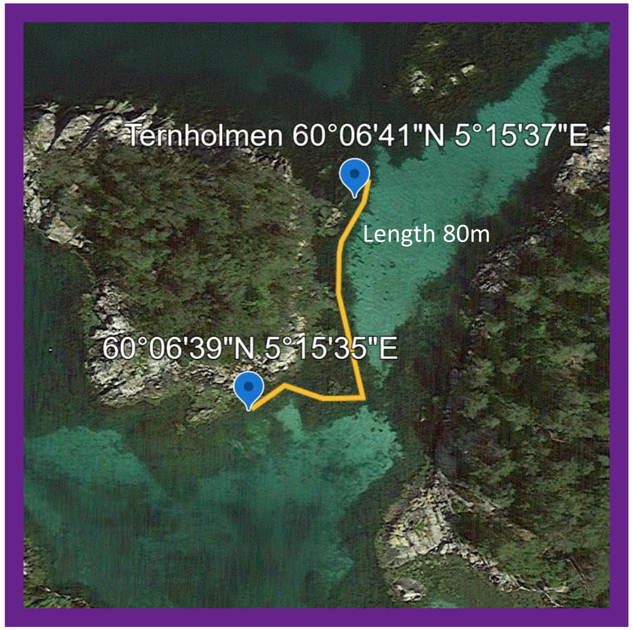


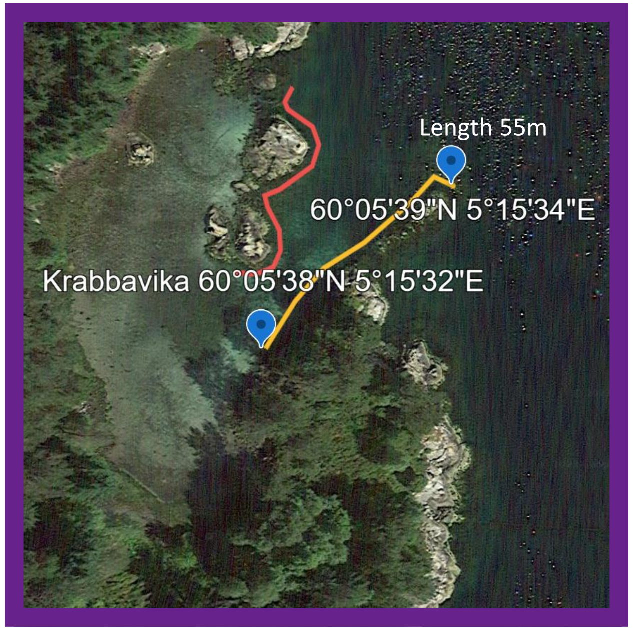


Images from Google Earth 2023

### Transect coordinates for the 5 sub-locations of the Hitra population.

Behavior and population census were recorded on transects indicated in yellow. Approximate transect length estimated from satellite image is given. Red lines indicate artificial nests lines.

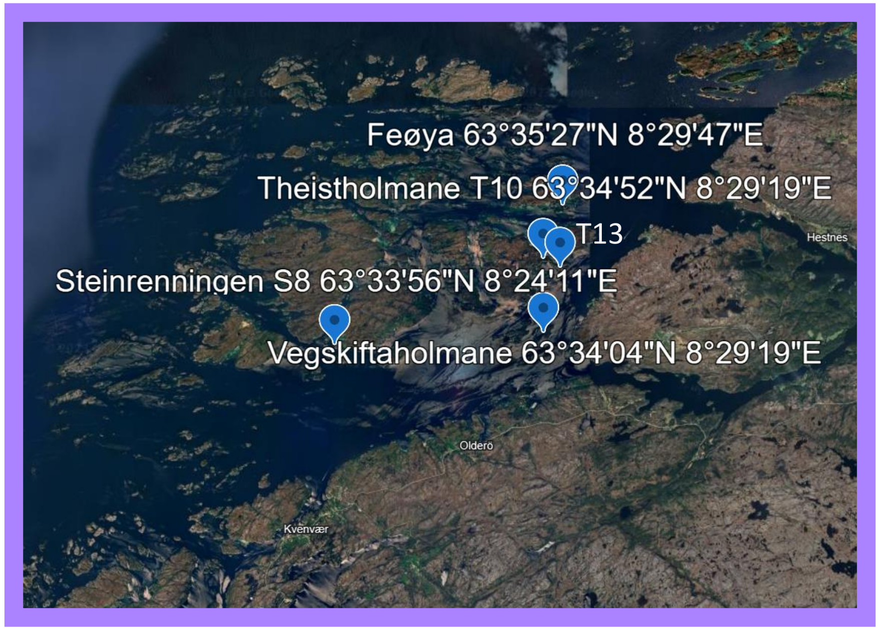


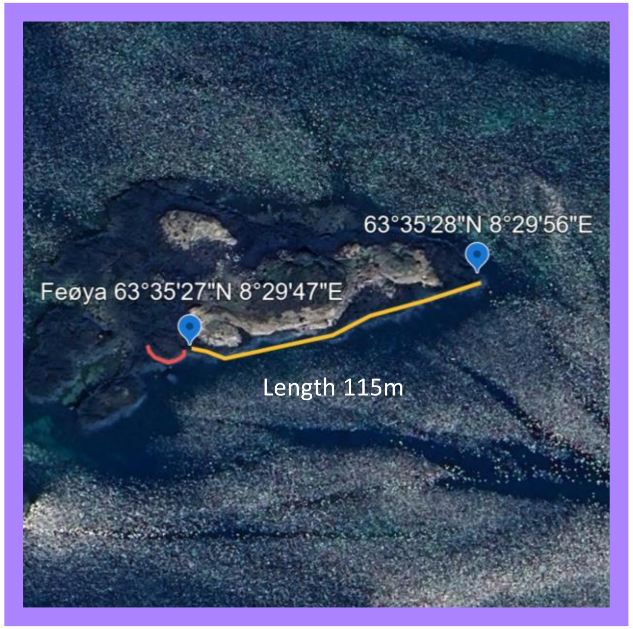


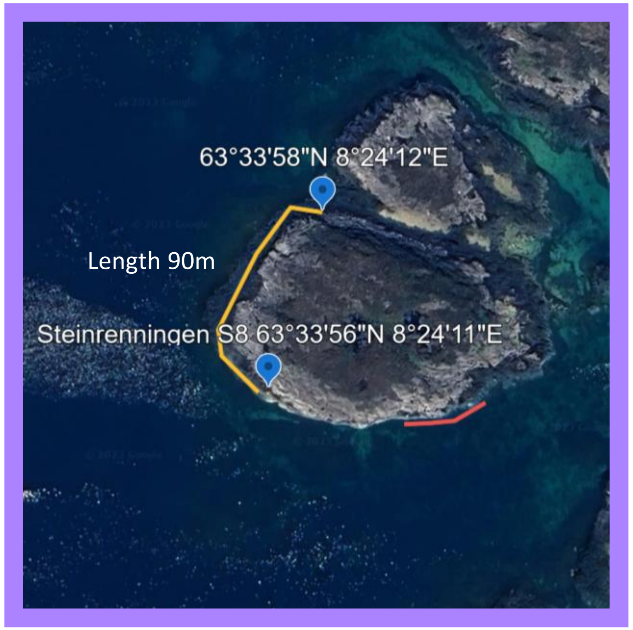


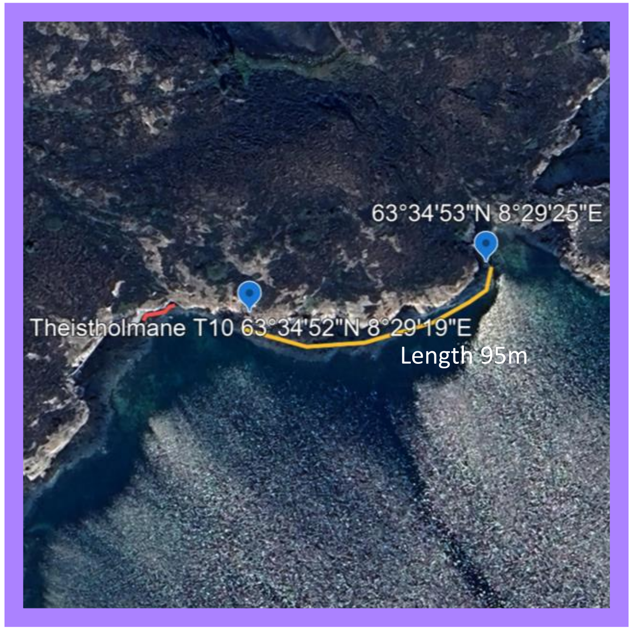


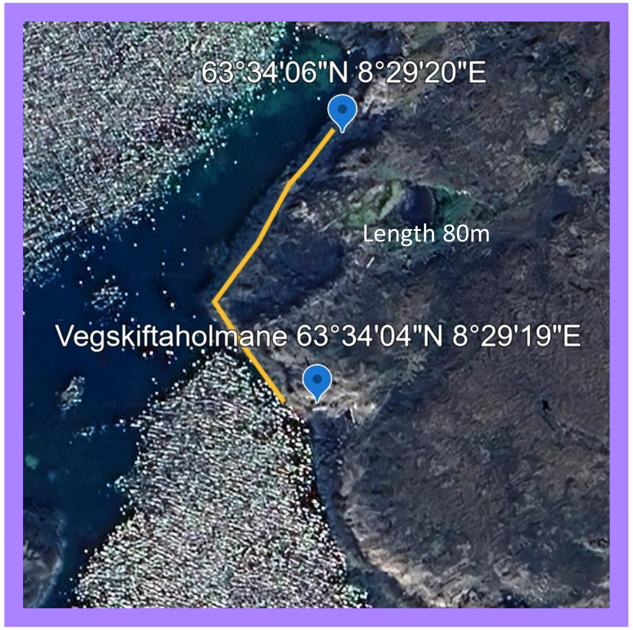


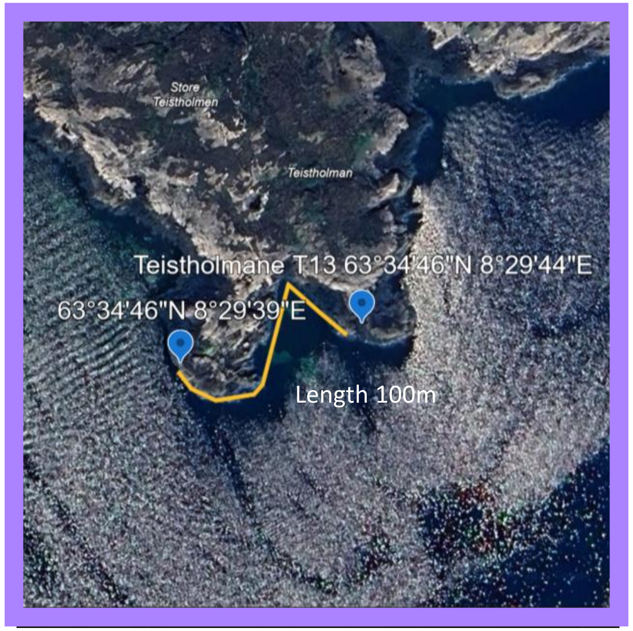


Images from Google Earth 2023

#### Transect coordinates for the 5 sub-locations of the Helligvær population.

Behavior and population census were recorded on transects indicated in yellow. Approximate transect length estimated from satellite image is given. Red lines indicate artificial nests lines.

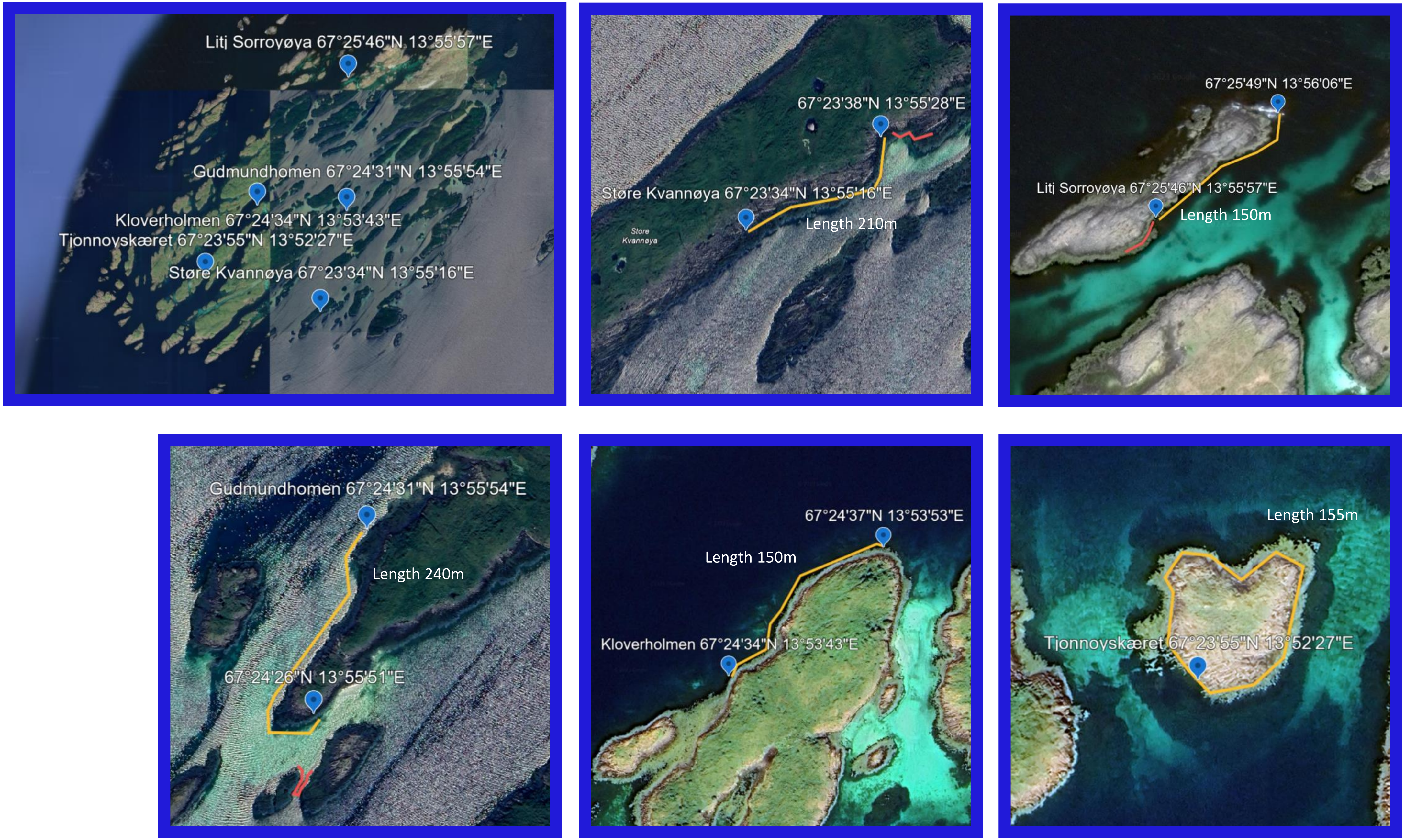


Images from Google Earth 2023

### Transect coordinates for the 5 sub-locations of the Ringstad population.

Behavior and population census were recorded on transects indicated in yellow. Approximate transect length estimated from satellite image is given. Red lines indicate artificial nests lines.

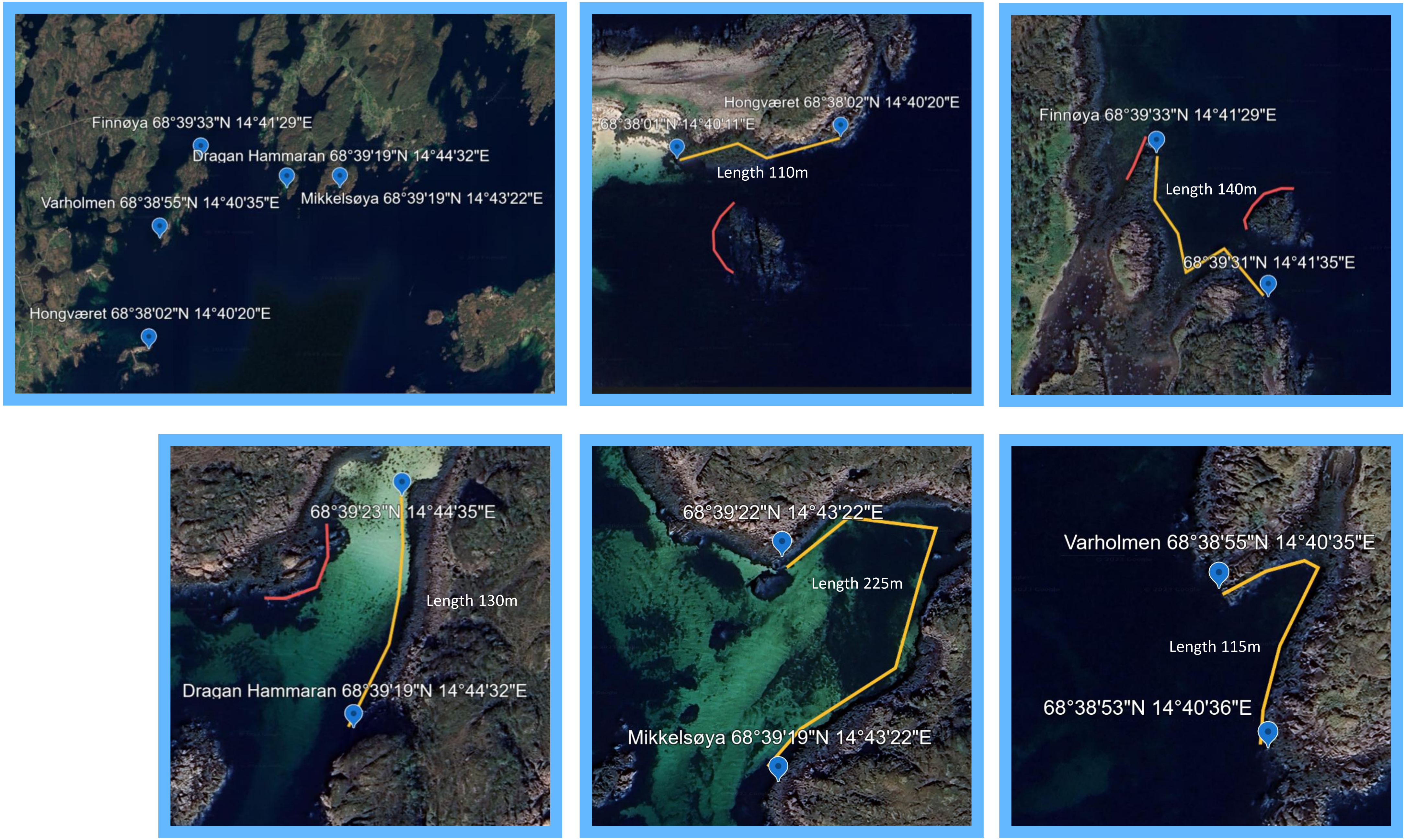


Images from Google Earth 2023
