## Supplementary Figures for "Sexual display behaviour follows consistent sex-specific reaction norms across latitude in response to operational sex-ratio"

### Supplementary S1: The six study populations, 5 sub-locations in each, and sampling schedule.

**a. The 6 populations**

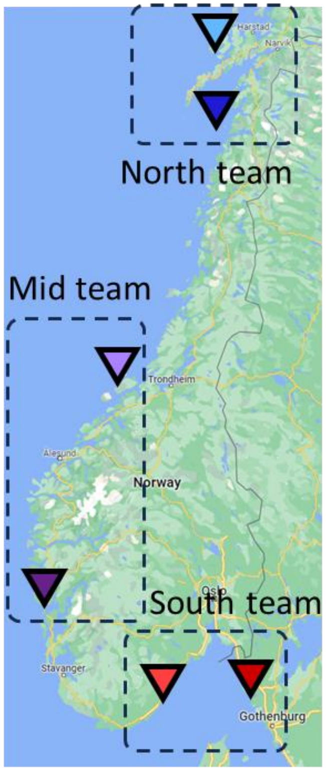

**b. The 5 sublocations for each population**

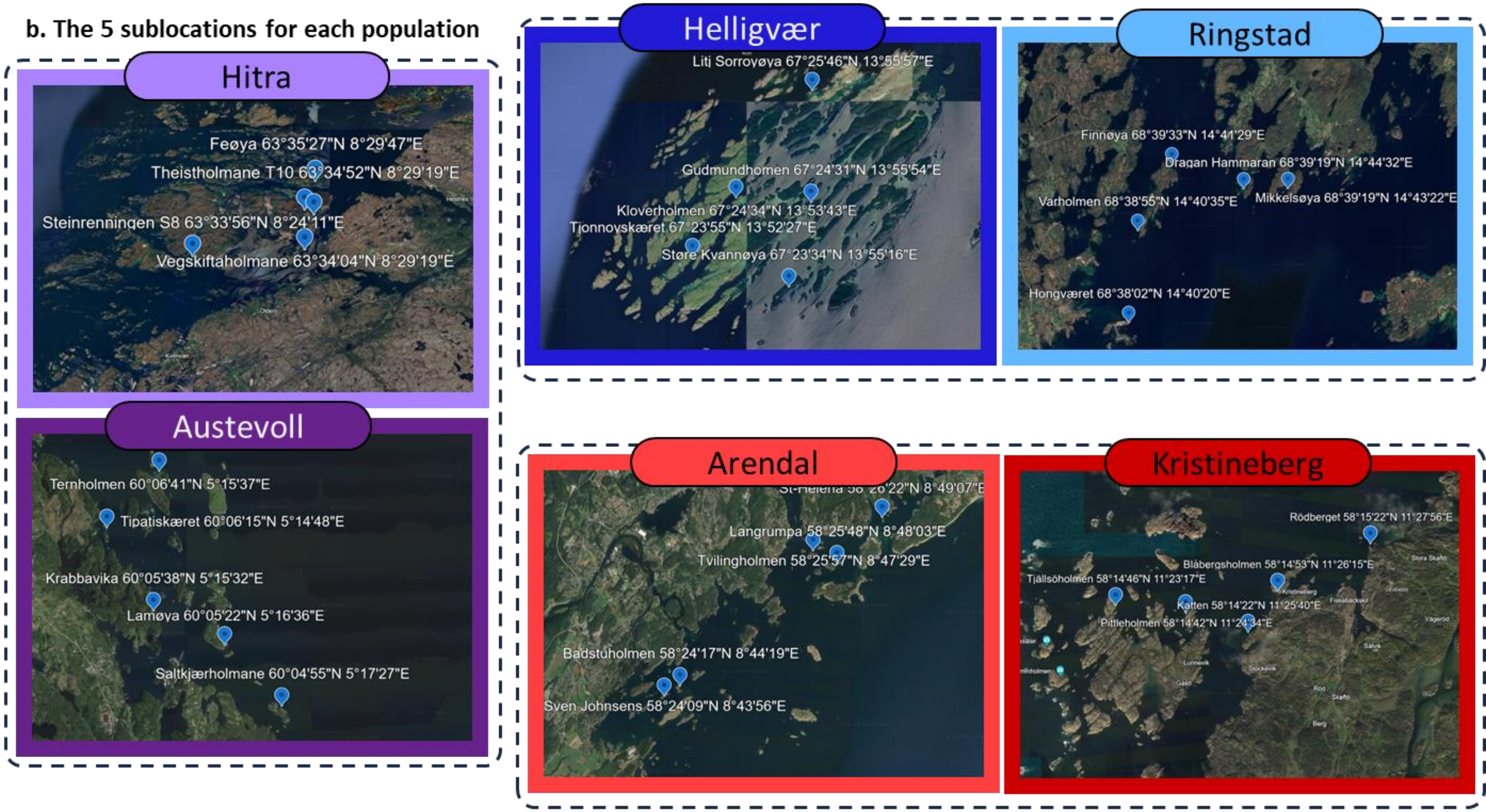

Maps from Google Earth 2023

**c. Timeline of the behavior and population census sampling for each sublocation**

Initials match the sublocation names in (b). Sampling was started later in northern locations to account for the expected delay in reproductive activity, matching delayed water temperature rise.

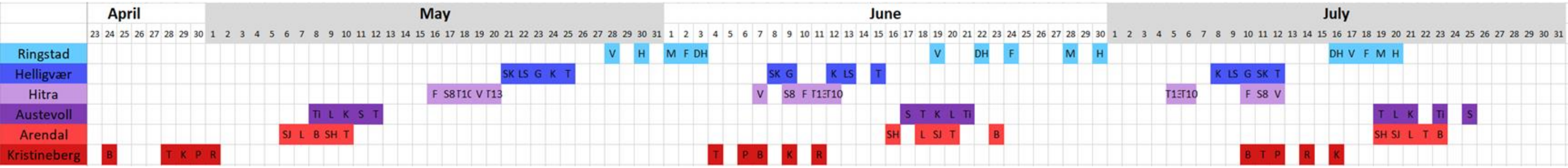

**BEHAVIOUR TRANSECT FORM** **DYNAMAR 2022**

| | | | |
|---|---|---|---|
| Area | Loc | Date | Obs |
| Waves cm | Sun Y/N | Rain Y/N | Temp |
| Start | Stop | Visibility | Sketch |

| **M to F** | Male action | | | Female response | | | |
|---|---|---|---|---|---|---|---|
| (M starts, one or both act) | **Fin** | # | | **Hook** | **Glow** | **Hook+Glo** | **No resp.** |
| | **Lead** | **Tow ?** | **Into N** | **Follow** | **Nest** | **Fol + Nest** | **No resp.** |

| **M & F** | Male action | | Female action | | | |
|---|---|---|---|---|---|---|
| (Both act, unclear or unknown who starts) | **Fin** | # | **Hook** | **Glow** | **Hook+Glo** | **No resp.** |
| | **Lead** | # | **Hook** | **Follow** | **Nest** | **No resp.** |

| **M to M** | Male Initiator | | Male Respondent | | |
|---|---|---|---|---|---|
| (One or both act) | **Fin** | # | **Fin** | **Retreat** | **No resp.** |
| | **Chase** | # | **Chase** | **Retreat** | **No resp.** |

| **F to M** | Female action | | Male response | | | | |
|---|---|---|---|---|---|---|---|
| (F starts, one or both act) | **Hook** | # | **Approach** | **Ap+Fin** | **AFLead** | **AFLNest** | **No resp** |
| | **Glow** | # | **Approach** | **Ap+Fin** | **AFLead** | **AFLNest** | **No resp** |
| | **Hook+Glo** | # | **Approach** | **Ap+Fin** | **AFLead** | **AFLNest** | **No resp** |

| **F to F** | Female Initiator | | | Female Respondent | | | |
|---|---|---|---|---|---|---|---|
| (one or both act) | **Hook** | **Glow** | **Hook+Glo** | **Hook** | **Glow** | **Hook+Glo** | **No resp.** |

COMMENTS

**Supplementary Figure S2. Behavior transect form for snorkeling observations**

M to F is male initiated behaviors
M & F is mutual courtship
M to M male agonistic behavior
F to M female initiated courtship
F to F female agonistic behavior
The form is printed on waterproof paper and carried along on a clipboard while snorkeling. When a behavior is observed, a tick mark is added in the first box of the corresponding panel, and the response of the other individual is ticked in the appropriate box.

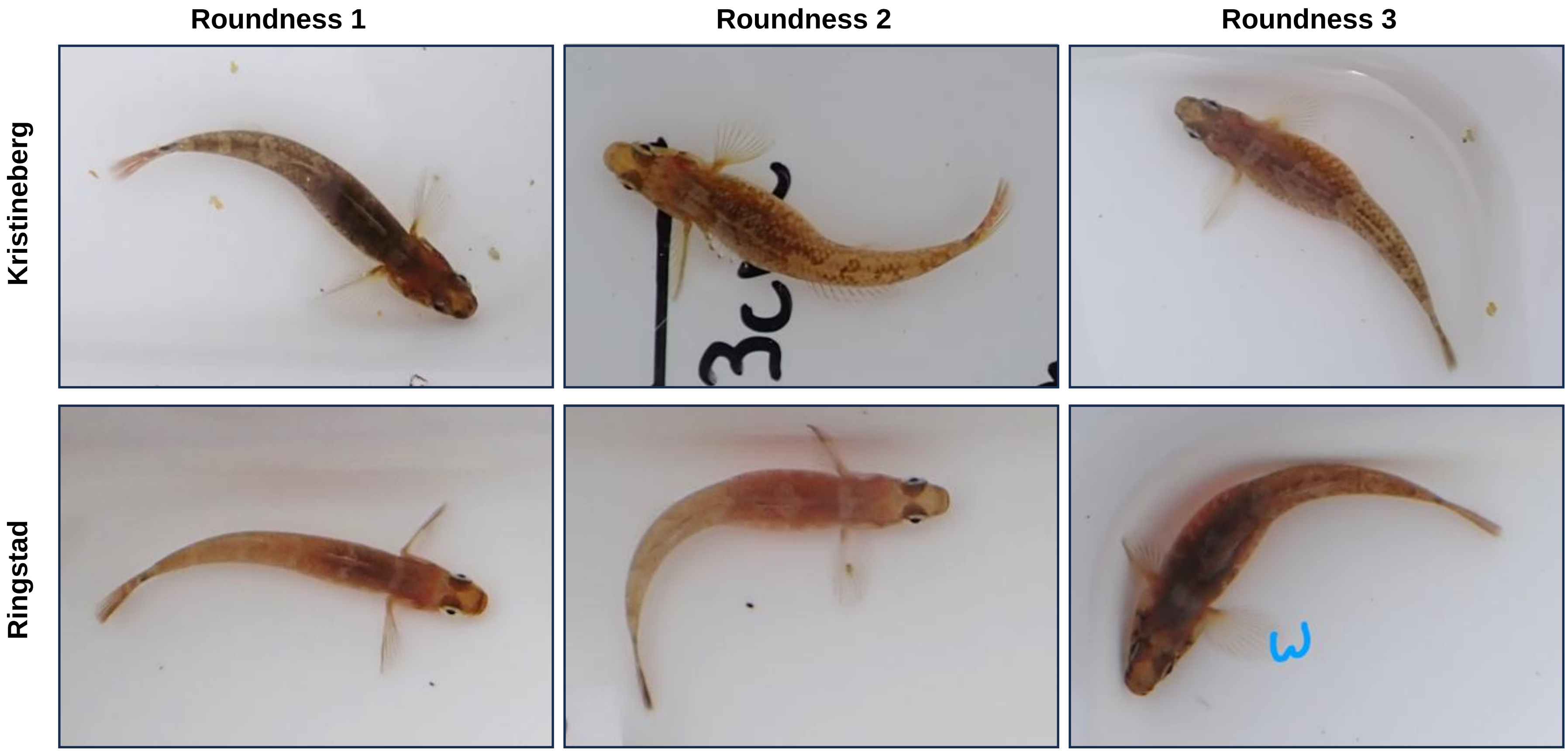

**Supplementary Figure S3.**
**Example of female roundness assessments for the Kristineberg and Ringstad populations.**
The roundness score is assessed subjectively by the field team leader. The three field team leaders trained in assessing roundness together prior to the start of the field season, until they achieved consistent assessments. Example pictures are given for the Southernmost and Northernmost populations.

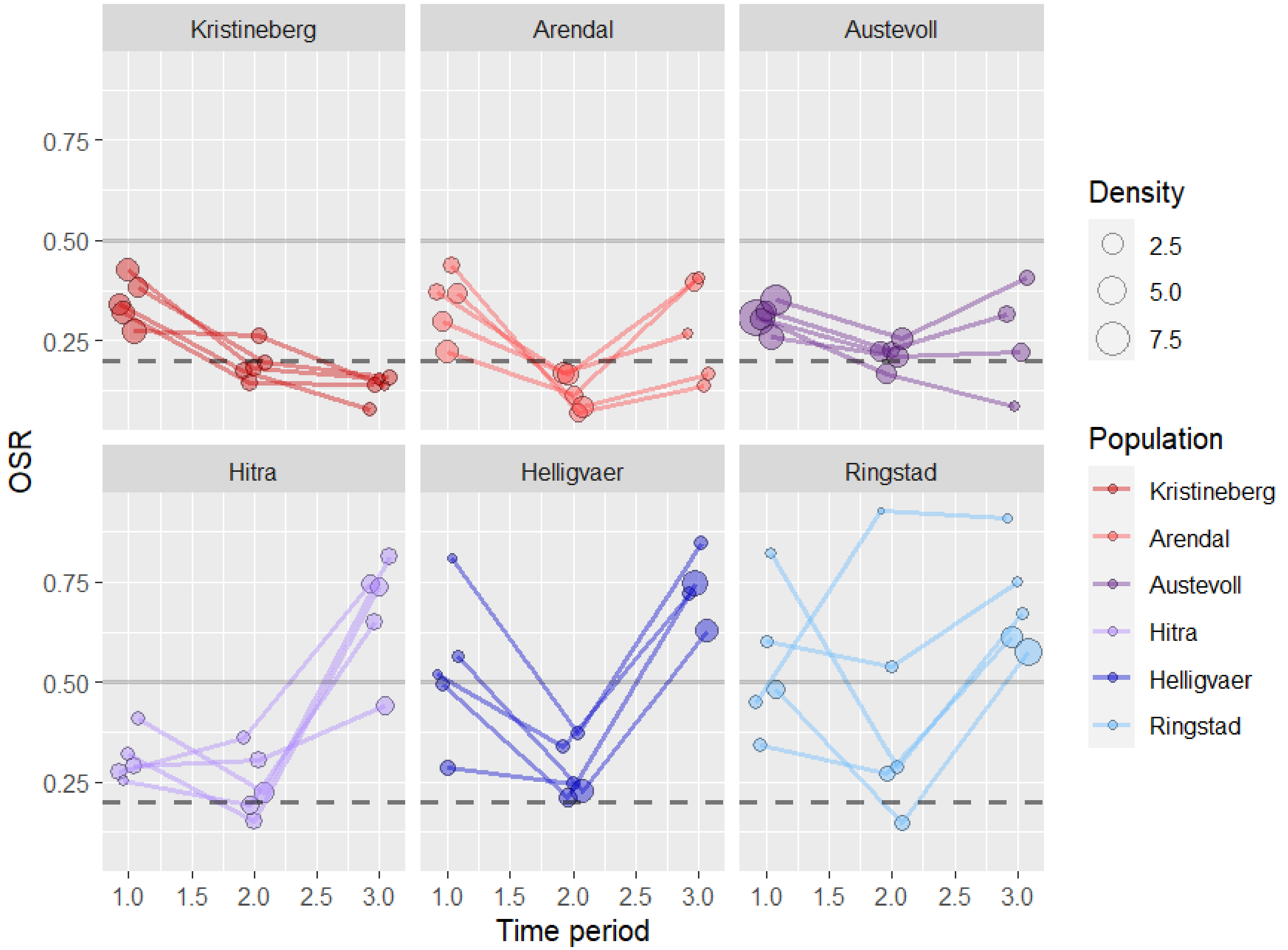

**Supplementary Figure S4. Social environment (OSR) for each study sublocation over time.** *This Supplementary  Figure is in complement to Figure 4 in the main manuscript. Each dot represents a population census event, with each study population replicated in 5 sublocations at each time period. The size of the dots represents population density (number of fish/meter), the colour refers to the population of origin. Lines are connecting the sublocactions across time. The solid grey horizontal line represents the switch point between males (OSR>0.5) or females (OSR<0.5) being more abundant in the mating pool. The dashed horizontal line represents the expected behavioural switching point with a when accounting for collateral investment (see Methods 4.4).*

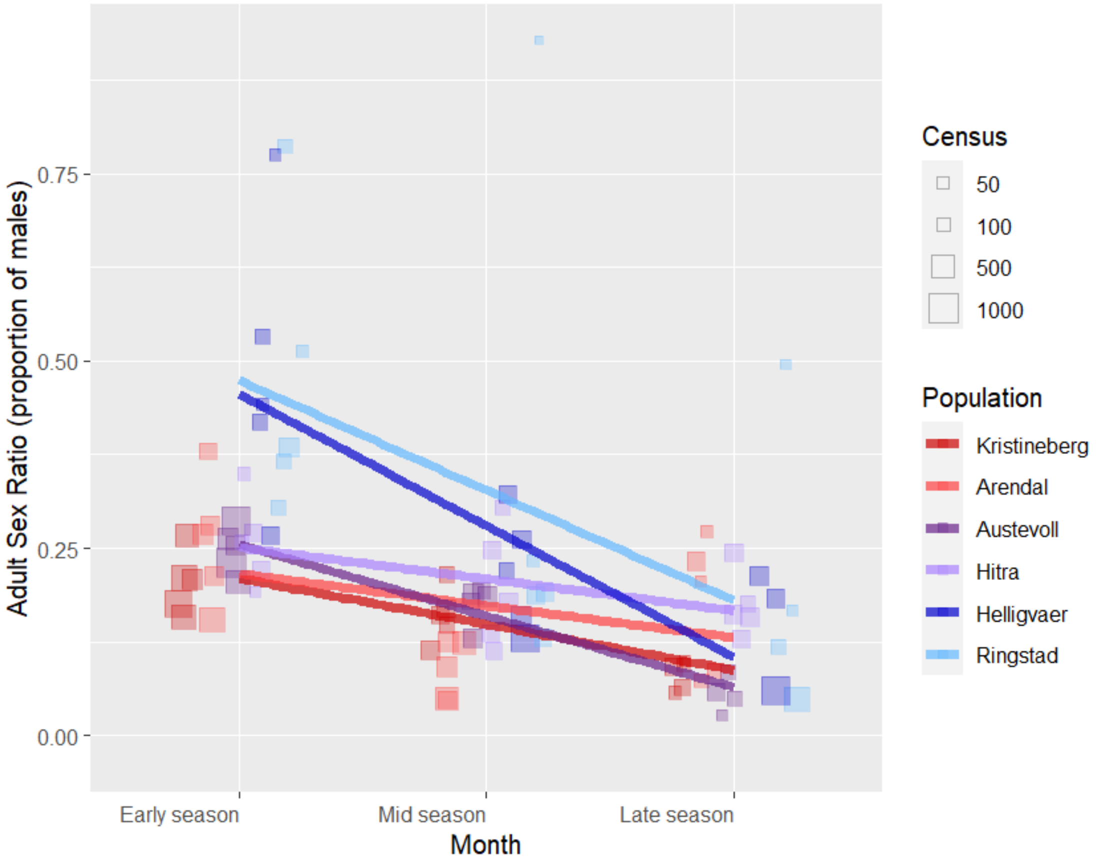

**Supplementary Figure S5. Adult Sex Ratio for each population and time period.**
*Each square dot represents a population census event with the size of the square being the total adult population count. ASR is calculated as the relative proportion of males in the adult population m/(m+f).*

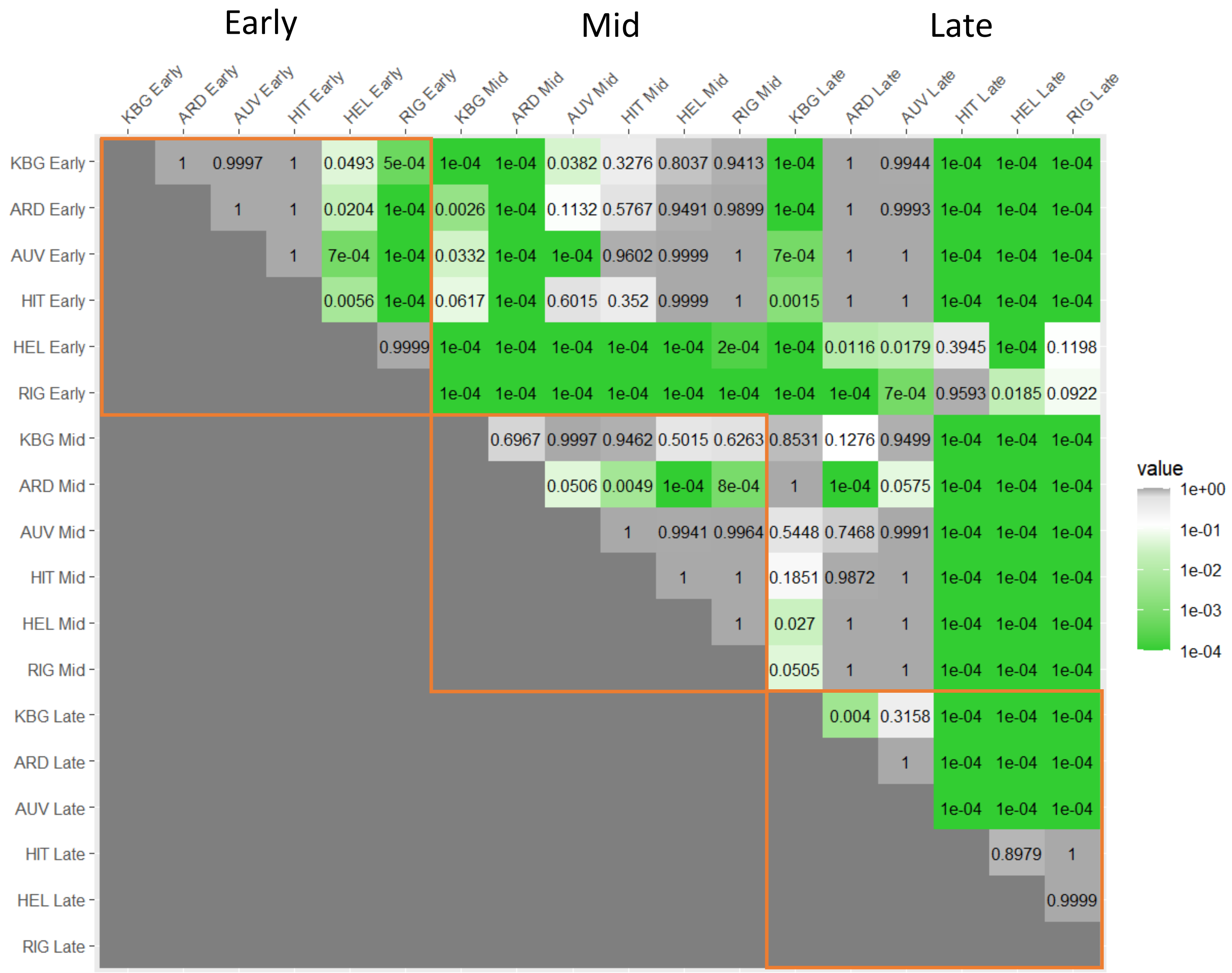

**Supplementary Figure S6. P- values of Tukey's post-hoc pairwise tests for Operational Sex Ratio (OSR) across populations and time periods.**

*The time periods are designated as Early, Mid- and Late season. Populations are in order from South to North (KBG=Kristineberg, ARD=Arendal, AUV=Austevoll, HIT=Hitra, HELL=Helligvær, RIG=Ringstad). The orange frames encompass within time period comparisons.*

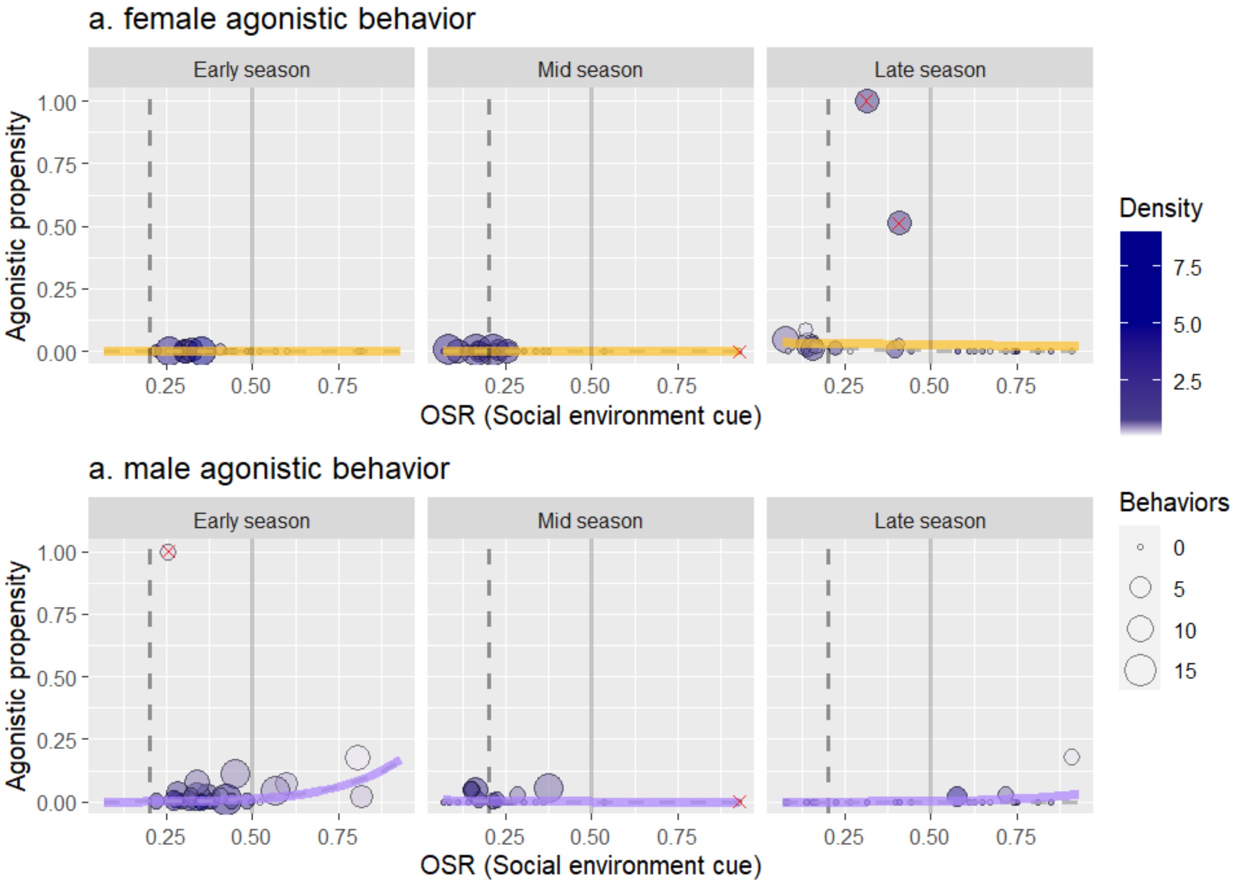

**Supplementary Figure S7. Sex-specific reaction norms for the propensity to perform same-sex agonistic behavior.** In support of main Figure 6, showing full y-axis range and outlier points. Each dot is a sublocation at a given sampling time (representing two consecutive observation transects), the size of the dots corresponds to the number of agonistic behaviours observed for the focal sex on that transect. The colour fill of the dots reflects the local density (individuals per meter of transect), where the white fill indicates low density (>0,75 fish/m) leading to higher stochasticity in the estimation of propensity. Agonistic propensity is estimated from the proportion of same-sex encounters leading to agonistic behaviour for the focal sex (see Methods 4.3). The solid grey vertical line represents the switching point between males (OSR>0.5) or females (OSR<0.5) being more abundant in the mating pool. The dashed vertical line represents the switching point between males or females being perceived as more abundant in the mating pool, taking into account the collateral investment (see Methods 4.4). Solid coloured trend lines represent predictions from a binomial generalised mixed-model fitted to the data. Data points marked with a red cross (5 in total) represent potential outliers, and the grey dashed trend lines represent model predictions when the outliers are excluded. For better visualisation, the y-axis was restricted to the 0-0.25 range, leading to the exclusion of 3 outlier points.
