## Supplementary Tables for "Sexual display behaviour follows consistent sex-specific reaction norms across latitude in response to operational sex-ratio"

**Supplementary table 1. Model comparison for courtship propensity data**

| Response | Outliers excluded | 3-way interaction | AIC | BIC | Skewness of residuals | Kurtosis of residuals |
|---|---|---|---|---|---|---|
| Courtship propensity | No | Yes | 1011 | 1053 | 0.61 | 4.39 |
| | | No | 1012 | 1040 | 0.56 | 4.39 |
| | Yes | Yes | 931.6 | 972.6 | 0.39 | 3.85 |
| | | No | 931.4 | 959.8 | 0.34 | 3.85 |

**Supplementary table 1. Model comparison for agonistic behavior data**

| Response | Outliers excluded | 3-way interaction | AIC | BIC | Skewness of residuals | Kurtosis of residuals |
|---|---|---|---|---|---|---|
| Agonistic probability | No | Yes | 681.7 | 722.9 | 1.33 | 7.26 |
| | Yes | | 558.2 | 599.2 | 0.46 | 3.38 |
| Agonistic propensity | No | Yes | 522.0 | 563.3 | 2.02 | 12.8 |
| | Yes | | 404.5 | 445.5 | 0.38 | 3.38 |
| | No | No | 519.2 | 547.7 | 2.00 | 12.7 |
| | Yes | | 400.5 | 428.9 | 0.37 | 3.51 |
